# IsoCon: Deciphering highly similar multigene family transcripts from Iso-Seq data

**DOI:** 10.1101/246066

**Authors:** Kristoffer Sahlin, Marta Tomaszkiewicz, Kateryna D. Makova, Paul Medvedev

**Author notes:** equal contribution (Kristoffer Sahlin, Marta Tomaszkiewicz).

## Abstract

A significant portion of genes in vertebrate genomes belongs to multigene families, with each family containing several gene copies whose presence/absence can be highly variable across individuals. For example, each Y chromosome ampliconic gene family harbors several nearly identical (up to 99.99%) gene copies. Existing *de novo* techniques for assaying the sequences of such highly-similar gene families fall short of reconstructing end to end transcripts with nucleotide-level precision or assigning them to their respective gene copies. We present IsoCon, a novel approach that combines experimental and computational techniques that leverage the power of long PacBio Iso-Seq reads to determine the full-length transcripts of highly similar multicopy gene families. IsoCon uses a cautiously iterative process to correct errors, followed by a statistical framework that allows it to distinguish errors from true variants with high precision. IsoCon outperforms existing methods for transcriptome analysis of Y ampliconic gene families in both simulated and real human data and is able to detect rare transcripts that differ by as little as one base pair from much more abundant transcripts. IsoCon has allowed us to detect an unprecedented number of novel isoforms, as well as to derive estimates on the number of gene copies in human Y ampliconic gene families.

## Introduction

A significant portion of genes in the human genome belongs to multigene families, with each family containing several gene copies that have arisen via duplication, *i.e*. duplicate gene copies^1 2 3 4 5 6^. Many of these duplicate genes have been associated with important human phenotypes, including a number of diseases ^7 8 9^. However, the annotation of multigene families remains incomplete even in the latest human assembly, especially due to unresolved segmental duplications with high sequence identity ^10,11^. Duplicate gene copies from the same family vary in sequence identity, with some of them being identical to each other. Additionally, copy numbers within families frequently differ among individuals ^1 2 3^. Furthermore, an estimated >90% of all multi-exon genes are alternatively spliced in humans ^12,13^, and different duplicate gene copies can vary in alternatively spliced forms (i.e. isoforms) produced.

These features make deciphering the end to end transcript sequences from duplicate genes and their various transcript isoforms a major challenge. The copy number of multigene families can be assayed using microarrays ^2^, quantitative PCR ^14^, droplet digital PCR ^15^, or DNA sequencing using Nanostring Technologies ^8^ or Illumina platforms ^3^. Sequences of individual exons that are only a few hundred nucleotides long can be obtained from individual reads of Illumina DNA or RNA-seq data^16^; however, the repetitive nature of duplicate gene copies complicates their *de novo* assemblies, and Illumina reads are often unable to phase variants across the length of the full transcript ^17 18^. Long Pacific Biosciences (PacBio) reads from the Iso-Seq protocol hold the potential to overcome this challenge by sequencing many transcripts end to end. This approach has been successfully applied to reveal several complex isoform structures resulting from alternative splicing events in, e.g., humans, plants, and fungi ^17,19,20^. None of these studies has simultaneously tackled the problems of deciphering isoform structure and of determining which gene copies they originated from.

While PacBio error rates have decreased, many errors remain hard to correct and remain a significant problem for downstream analyses of Iso-Seq data ^21,22,18^. This is especially the case for transcripts from gene families with high sequence identity, where teasing out errors from true variants is difficult. The use of a reference genome ^23,24,25,26,27^ for correction is not effective in such situations, where the variability of gene copies might not be reliably captured by the reference. ICE^17^, a part of PacBio’s bioinformatic pipeline to process Iso-Seq data, is the standard tool employed to correct sequencing errors without using the reference. Though ICE has been utilized in several projects ^28 29 28^, it has been shown to generate a large number of redundant transcripts ^17,20,30^. Moreover, ICE “is not currently customized to work for differentiating highly complex gene families from polyploid species where differences are mostly SNP-based.” ^31^ An alternate approach -- to use Illumina reads to correct errors in PacBio reads ^24,27,25,32^ -- is similarly unable to correct most errors (as we demonstrate in this paper) and is also biased by low Illumina read depth in GC-rich regions ^33^.

Some approaches were proposed to decipher and error correct PacBio reads from transcripts with high sequence identity, but none is broadly applicable to determining the sequences from high-sequence-identity multigene families without relying on the reference genome. Classification ^34^ or construction ^35^ of allele-specific transcripts with Iso-Seq have been described, but these approaches require a reference and can only separate two alleles of a single gene. Genotyping approaches for multigene families have also been proposed ^36^, but they require prior knowledge of the isoform sequences. A *de novo* approach for clustering highly similar isoforms is described in ^37^, but no implementation is provided. The problem is also related to that of viral phasing ^38^, but the techniques developed are not directly applicable to human multigene families.

Another consideration is the relatively high cost of PacBio. The number of reads required to recover gene families whose expression is dwarfed by super-prevalent mRNA classes can be prohibitive. A targeted sequencing approach can be effective at reducing the necessary amount of sequencing, where RT-PCR primer pairs are designed to pull out transcripts of the gene family of interest ^39^. This approach results in sequencing depths high enough to capture most transcripts and perform downstream error correction.

To address many of these limitations, we develop IsoCon, a *de novo* algorithm for error-correcting and removing redundancy of PacBio circular consensus sequence (CCS) reads generated from targeted sequencing with the Iso-Seq protocol. Our algorithm allows one to decipher isoform sequences down to the nucleotide level and hypothesize how they are assigned to individual, highly similar gene copies of multigene families. IsoCon uses a cautiously iterative process to correct obvious errors, without overcorrecting rare variants. Its statistical framework is designed to leverage the power of long reads to link variants across the transcript. Furthermore, IsoCon statistically integrates the large variability in read quality, which decreases as the transcript gets longer. Using simulated data, we demonstrate that IsoCon has substantially higher precision and recall than ICE ^17^ across a wide range of sequencing depths, as well as of transcript lengths, similarities and abundance levels. IsoCon is able to capture transcripts that differ by only one nucleotide in sequence and by three orders of magnitude in abundance.

We apply IsoCon to the study of Y chromosome ampliconic gene families, where the inability to study separate gene copies and their respective transcripts has limited our understanding of the evolution of the primate Y chromosome and the causes of male infertility disorders for which these genes are crucial ^40 41 42 43^. Y chromosome ampliconic gene families represent a particularly interesting and challenging case to decipher, because each of them contains several nearly identical (up to 99.99%) copies ^44 45^ with a potentially varying number of isoforms. We use a targeted design to isolate and sequence all nine Y chromosome ampliconic gene families from the testes of two men. Our validation shows that IsoCon drastically increases precision compared to both ICE and Illumina-based error correction with proovread ^46^ and significantly higher recall than ICE. We show that IsoCon can detect rare transcripts that differ by as little as one base pair from dominant isoforms that have two orders of magnitude higher abundance. Using IsoCon’s predicted transcripts, we are able to capture an unprecedented number of isoforms that are absent from existing databases. We are further able to separate transcripts into putative gene copies and derive copy-specific exon sequences and splice variants.

IsoCon is open-source and freely available at https://github.com/ksahlin/IsoCon. All analyses are reproducible via scripts and snakemake ^47^ workflows at https://github.com/ksahlin/IsoCon_Eval.

## Methods

### IsoCon

#### Overview

The input to the IsoCon algorithm is a collection of PacBio circular consensus sequence (CSS) reads and their base quality predictions. IsoCon assumes the reads have been pre-processed to remove primers, barcodes, and reads that are chimeras or do not span the whole transcript. The pre-processing step separates the reads according to the primer pairs used to amplify individual gene families, and IsoCon is run separately on each gene family. The output of IsoCon is a set of transcripts which are the result of error-correcting the reads and reporting each distinct read.

IsoCon consists of two main steps: (i) an iterative clustering algorithm to error-correct the reads and identify candidate transcripts, and (ii) iterative removal of statistically insignificant candidates.

The clustering/correction step partitions the reads into clusters, where reads that are similar group together into one cluster. A multiple alignment and a consensus sequence is computed for each cluster. The reads in each cluster are then partially error-corrected to the cluster’s consensus sequence; to avoid removing true variants, only half of the potentially erroneous columns are corrected. Then, the process iterates -- the modified reads are repartitioned into potentially different clusters and corrected again. This process is repeated until no more differences are found within any cluster, and the distinct sequences remaining are referred to as candidate transcripts (or simply candidates).

The clustering/correction step is designed to be sensitive and is therefore followed by the second step, which removes candidate transcripts that are not sufficiently supported by the original (non-corrected) reads. Initially, the original reads are assigned to one of their closest matching candidates. Then, evaluating all pairs of close candidates, for every pair we check whether there is sufficient evidence that their assigned reads did not in fact originate from the same transcript. To do this, we take two candidates and their set of variant positions and formulate a hypothesis test to infer how likely it is that the reads supporting these variants are due to sequencing errors. Since a candidate can be involved in many pairwise tests, it is assigned the least significant p-value from all pairwise tests performed. After all pairs of candidates have been tested, a fraction of non-significant candidates will be removed. The second step of IsoCon is then iterated -- the original reads are assigned to the remaining candidates, which are then statistically tested. This continues until all remaining candidates are significant. The remaining candidates are then output as the predicted transcripts.

#### Clustering/correction step

First, we need to define the concept of closest neighbors and the nearest neighbor graph. Let *dist*(*x,y*,) denote the edit distance between two strings *x* and *y*. Let *S* be a multi-set of strings. Given a string, *x* and *y* Let *S* be a multi-set of strings. Given a string *x*, we say that a *y* ∈ *S* is *closest neighbor of x* in *S* if 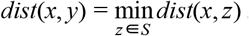. That is, *y* has the smallest distance to *x* in *S*. The nearest neighbor graph of *S* is a directed graph where the vertices are the strings of *S*, and there is an edge from *x* to *y* if and only if *y* is a closest neighbor of *x* but is not *x*.

There are two phases to the clustering/correction step – the partitioning phase and the correction phase – and we iterate between the phases. In the partitioning phase, we first partition the reads into clusters, with each cluster having exactly one read denoted as the *center*. The idea is that each partition contains a putative set of reads that came from the same transcript and the center is the read whose sequence is most similar to that of the transcript. To partition, we first build a graph *G* which, initially, is identical to the nearest neighbor graph built from the reads. Next, we identify a read *x* in *G* with the highest number of vertices that can reach *x*. We create a new cluster with *x* as the center and containing all reads in *G* that have a path to *x*, including *x* itself. Next, we remove the elements of the new cluster, along with their incident edges, from *G*. Then, we iterate on the newly modified *G*: identifying the vertex with the highest in-degree and creating a cluster centered around it. The full pseudo-code is given in the PartitionStrings algorithm in Fig. S1.

The resulting partition has the property that each string has one of its closest neighbors (not including itself) in its cluster. This closest neighbor may be the center but does not have to be. Thus, a cluster may contain many strings which are closest neighbors of others but only one of them is denoted as the center.

The correction phase works independently with each cluster of reads and its corresponding center. We first create pairwise alignments from each read to the center. We then create a multi-alignment matrix *A* using the standard progressive alignment method ^48^. Each entry in *A* is either a nucleotide or the gap character, and each row corresponds to a read. We obtain the consensus of *A* by taking the most frequent character in each column. Every cell in *A* can then be characterized as one of four states with respect to the consensus: a substitution, insertion, deletion, or match. Given a column *j* and a state *t*, we define 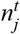 as the number of positions in column *J* that have state *t*. Similarly, let *n_t_* denote the total number of cells with state *A* in *A*. The *support* for a state *t* in column *j* is defined as 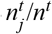, and the support of a cell in *A* is the support for that cell’s state in that cell’s column. In each read, we identify the variant positions (*i.e*. whose state is substitution, insertion, or deletion) and select half of these position that have the lowest support. Then, for each of these positions, we correct it to the the most frequent character in the column; but, if the most frequent character does not exist or is not unique, then no correction is made.

IsoCon’s clustering/correction step combines the partitioning and correction phases in the following way. Initially, we partition the set of reads and correct each cluster. A cluster is said to have converged if all its strings are identical. As long as at least one cluster is not converged, we repeat the partitioning and correction phases. To ensure that eventually all clusters converge, we heuristically undo the correction of a string if, after correction, it has a higher edit distance to the center than it had to its center in the previous iteration, if the string alternates between partitions in a cyclic fashion, or the same set of strings repeatedly get assigned to the same partition where they differ only at positions where the most frequent character is not defined. Finally, after all the partitions have converged, we designate their centers as *candidate transcripts* and pass to the candidate filtering step of IsoCon. The full pseudo-code for this step is given in the ClusterCorrect routine in Fig. S1.

#### Candidate filtering step

IsoCon’s second step takes as input a collection of reads *X* and a set of candidate transcripts *C* = {*c_1_*,… *c_*l*_*}.The first step is to assign reads to candidates, such that one read is assigned to exactly one of its closest neighbor candidates in *C*. Because a read may have several closest neighbor candidates in *C*, there are many possible assignments. For our purposes, we use the following iterative greedy algorithm. For each read *x* ∈ *X*, we identify its closest neighbor candidates in *C*. Next, we select a candidate *c* ∈ *C* that is a closest neighbor of the most reads in *X*. We assign all these reads to *c*, and remove *c* from *C* and all the assigned reads from *X*. We then repeat the process, using the reduced *X* and *C*, until all reads have been assigned.

Now, we have an assignment of reads to the candidates. We denote by *X_t_* the reads that are assigned to candidate *c_t_*. We check for evidence to support that *c_i_* is a true candidate as follows. We consider the candidates who are the closest neighbors of *c_t_* in *C* – {*c*_*i*_}. Next, for each closest neighbor candidate *c_j_*, we calculate the confidence that the reads in *X_i_* and in *X_j_* originated from *c_j_, i.e*. c_*i*_ is not a true candidate. The significance value calculation is given in the next section. We compute ρ_*i*_, the least significant value, amongst all *c_j_*. We limit our comparisons of *c_i_* to only its closest neighbor candidates because it keeps our algorithm efficient and it is unlikely that comparison against other more dissimilar candidates would increase ρ_*i*_.

Then, we identify the candidates with ρ_*i*_ greater than a significance threshold α. This α is a parameter to our algorithm, set by default to 0.01. These candidates are then removed from the candidate list *C*. Given a parameter τ, if there are more than τ candidates with significance value over α, we only remove the top τ candidates with the highest values. The candidate filtering step of the algorithm then iterates: we again assign reads to candidates and identify candidates with insufficient support. The algorithm stops when there are no longer any candidates with ρ_*i*_ above the significance threshold. The pseudo-code for this algorithm, together with all of IsoCon, is given in Fig. S1.

#### Statistical test

We are given two candidate transcripts *c* and *d* and sets of reads *X_c_* and *X_d_* that have been assigned to them. We use *x_i_* ∈ *X_c_* ∪ *X_d_* to denote each read and let *n* be the number of reads in *X_c_* ∪ *X_d_*.calculate pairwise alignments from *X_c_* ∪ *X_d_* ∪ {*c*}to *d*. Next, we progressively combine these alignments to construct a multi-alignment matrix *A*. Each entry in *A* corresponds to either a nucleotide or the gap character. Let *V* be the index of *A* the columns of*A* where *c* and *d* do not agree. We refer to these positions as *variant positions*.

Let *A_i_*,*j* denote the character in column *j* of row *i* of *A*. Rows 1 ≤ *i* ≤ *n* corresponds to reads *x_i_*. while row *n* + 1 corresponds to *c*. For 1 ≤ *i* ≤ *n*, we define a binary variable *S_i_* that is equal to 1 if and only if *A_i_*,*j* = *A_n_*+1,*j* for all *j* ∈ *V*. That is *S_i_*, is 1 if and only if read *x_i_* supports all the variants *V*, *i.e*. has the same characters as *c* at positions*V* in *A*. We make the following assumptions:

1. *d*, acting as the reference sequence in this test, is error-free.
2. A nucleotide in a read at a position that is not in *V* and differs from the corresponding nucleotide in *d* is due to a sequencing error. In other words, at position where *c* and *d* agree, then they cannot be both wrong.
3. The probabilities of an error at two different positions in a read are independent.
4. *S_i_* and *S_i_* are independent random variables for all *i* ≠ *i* ′.

Our null-hypothesis is that the variant positions in *A* are due to sequencing errors in *X*. To derive the distribution of *S_i_* under the null-hypothesis, we first need a probability, denoted by *p_ij_*, that position *j* in read *i* is due to an error. This can largely be obtained from the Phred quality scores in the reads (see *i* section Sup. Note A for details). Under assumption 3 we have that *S_i_* follows a Bernoulli distribution with a mean 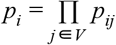

A relevant test statistic under the null-hypothesis is a quantity that models the strength (or significance) of support for variants *V*. We would like to only count reads that fully support all the variants, *i.e*.reads *x_i_* with *s_i_* = 1 (*s_i_* denotes the observed value of *S_i_*). These reads may have errors in non-variant locations, but, at the variant locations, they must agree with *c*. For each such read, we would like to weigh its contribution by the inverse of the probability that all the characters at the variant locations are due to sequencing errors. Intuitively, a read with a high base-quality should count as more evidence than a read with a low base-quality. Taking these considerations together, we define our test statistic as 

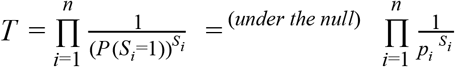

Notice that *p_i_* decreases with the amount of variants in V and with higher base quality scores; therefore, *T* is designed to leverage linked variants across the transcript, in the sense that less reads are required to support a transcript when the transcript has more variants. Moreover, *p_i_* decreases for reads with higher CCS base quality at variant positions, meaning less reads are needed to support a transcript, if they have higher quality. We observed that base quality values in the CCS was highly variable and depends on (i) the number of number of passes in a CCS read, (ii) the mono-nucleotide length and (iii) the sequenced base, with C and G having lower qualities associated with them (Fig. S2)

We let *t* be the observed value of this statistic and we refer to it as the *weighted support*. Given *t*, we calculate a significance value as *P* (*T* ≥*t*). We use a one-sided test as we are only interested in significance values of equal or higher weighted support for *V*. We are not aware of a closed form distribution of *T* under the null-hypothesis, and a brute-force approach to calculating *P* (*T* ≥ *t*) would be infeasible. However, we can make use of the following Theorem from ^49^, which gives a closed formula upper-bound on the distribution of a sum of Bernoulli random variables:

##### Theorem

Let *a*_1_,…, *a_n_* be reals in (0,1] and *Z*_1_,…, *Z_n_* be Bernoulli random trials. Let *Z* = ∑ *a_i_ Z_i_* and δ > 0 μ = *E* (*Z*) ≥ 0, then 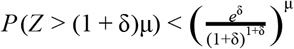

In order to apply the Theorem to T, we must first make a log transformation to convert the product into a sum, and then normalize *T* so that coefficients lie in (1,0] as needed. 

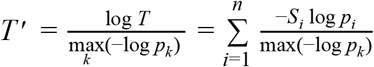

The expected value is 

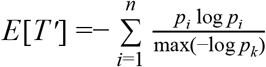

Note that under this transformation, P(*T*′ ≥ *t*′) = *P* (*T* ≥ *t*), as the logarithm function is strictly monotone and the normalization using the maximum is constant. Let μ= *E*(*T*′) and μ=*t*′/μ.We can then apply the Theorem to obtain the bound 

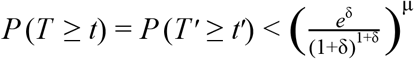

We use this upper bound as the significance value. Note that the Theorem only applies for δ> 0. If*t*′ ≤ μ,then this is not the case. However, it implies that the observed weighted support is below the expected support, under the null-hypothesis. Such values are clearly insignificant, and our software defaults to a value of 0.5.

## Experimental methods

Poly(A) RNA was isolated from testis RNA of two Caucasian men (IDs: CR560016, age 59, sample 1; CR561118, age 79, sample 2; Origene) using Poly(A) Purist MAG kit (Thermo Fisher Scientific). 500 ng of poly(A) RNA per each sample, along with 1 ug of control liver total RNA (used for control), were used to generate double-stranded DNA using SMARTer PCR cDNA Synthesis Kit (Clontech). PCR cycle optimization of cDNA amplification reaction using the Clontech primer was performed and 12 cycles were determined to be optimal for the large-scale PCR amplification. For each of nine ampliconic gene families, we designed a pair of RT-PCR primers with one primer located in the first, and the other primer located in the last, coding exon (Table S1). For one of these gene families (*CDY*), an additional primer pair was designed to capture transcripts originating from all gene copies (Fig. S3). One of the two unique PacBio barcodes was added to the primers in order to distinguish RT-PCR products between the two men. Next, RT-PCR products from these two individuals were separated into two equimolar pools according to the expected transcript sizes (<1 kb and 1–2 kb; Table S1) and purified using AMPure XP beads (Beckman Coulter, Inc., USA). Each of the two RT-PCR pools was then used to construct a separate PacBio Iso-Seq library that was sequenced on RSII (P6-C4 chemistry) using one SMRT cell per library. Therefore, a total of two SMRT cells were sequenced.

Additionally, we sequenced the same RT-PCR products with Illumina technology. We constructed separate Nextera XT library (with a unique pair of indices) for each primer pair-sample combination. A total of nine gene families were analyzed with 10 primer pairs (as mentioned above, one gene family, *CDY*, was analyzed with two primer pairs). Therefore, 10 primer pairs x 2 individuals = 20 libraries were constructed. These libraries were normalized, pooled in equimolar ratio, and sequenced on a MiSeq instrument using one MiSeq Reagent Nano Kit, v2 (250x250 paired-end sequencing).

## Results

We evaluated IsoCon on several simulated datasets and a biological dataset of targeted sequencing from nine Y chromosome ampliconic gene families.

### Simulated data

We generated synthetic gene families using three reference genes as the starting sequence for our simulation: *TSPY*, *HSFY*, and *DAZ*. We chose these because they reflect the spectrum of length, exon number, and complexity, that is characteristic of Y ampliconic gene families (Table 1). *DAZ* is the hardest case, since it has a highly repetitive exon structure^50^. Gene length is also important, since longer transcripts result in fewer passes of the polymerase during sequencing and, hence, a higher error rate of CCS reads. We simulated coverage levels in a range consistent with what we later observed in real data.

**Table 1.** Gene sequences (taken from the corresponding reference gene in Ensembl) and their corresponding PacBio CCS error rates used for simulation. We simulated multigene families by using these gene sequences at the root. We refer to the resulting gene families by using the name of the reference gene, sometimes dropping the last modifier (*i.e. TSPY instead of TSPY13P*).

| Reference gene | Sum of exon lengths (nt) | Number of exons | Simulated error profile |  |  |
| --- | --- | --- | --- | --- | --- |
|  |  |  | Overall error rate | Median errors per transcript | Median n. of Passes |
| *TSPY13P* | 914 | 6 | 0.5% | 2 | 18 |
| *HSFY2* | 2,668 | 6 | 2.6% | 41 | 6 |
| *DAZ2* | 5,904 | 28 | 6.1% | 276 | 2 |

Our main simulation focused on two scenarios. The first one (Fig. 1) reflects a typical biological scenario. For each of the three gene families, we simulated several gene copies and, for each copy, we simulated various isoforms by skipping different exons. There were a total of 30 simulated isoforms per family, with relative abundances randomly assigned and ranging from 0.1% to 15%. We generate three such replicates by varying the mutation rate used to generate duplicate gene copies. (Note that here for simplicity we model mutation only, although other processes, e.g., gene conversion, are known to influence evolution of duplicate gene copies ^51^). The second simulation (Fig. S4) is similar to the first, but, in order to tease out the effect of mutation from that of exon skipping, we do not simulate isoforms. For each gene family, there were a total of eight gene copies and eight transcripts (one per gene copy) simulated, with varying sequence identity (Fig. S5) and with relative abundances ranging from 0.4% to 50%. We also repeated these two simulations but kept the isoform abundance constant (Figs. S6 and S7). See Sup. Note B for a complete description of our simulation.

**Figure 1.**
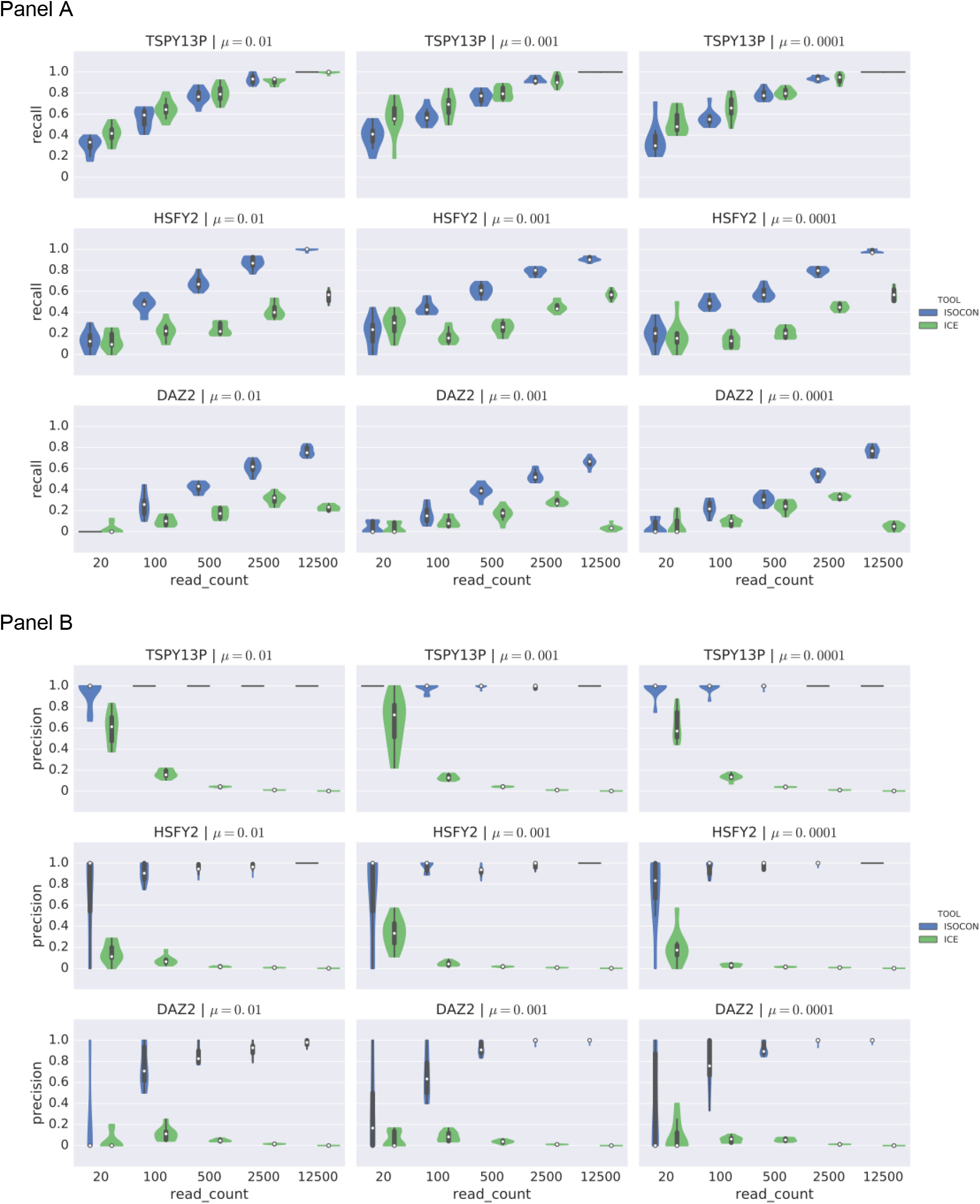
Violin plots showing the recall (panel A) and precision (panel B) of IsoCon and ICE on simulated families of transcripts with different exon structure and unequal abundance rates.In each panel, the rows correspond to different families and, hence, different error rates. The shortest gene family *TSPY*, labeled by the name of the reference gene copy used to generate it, *TSPY13P*), with correspondingly lower read error rates, is shown in the top panel rows while the longest gene family (*DAZ*, labeled as *DAZ2*), with correspondingly higher read error rates, is shown in the bottom rows. The columns correspond to different mutation rates (used in simulating the gene copies (see Sup. Note B).(μ) A lower mutation rate implies more similar gene copies. Each plot shows results for a total of 30 isoforms with abundances randomly assigned and ranging from 0.1% to 15%. Within each plot, the x-axis corresponds to the number of simulated reads, while the y-axis shows the recall/precision of the methods. Each “violin” is generated using ten simulated sequencing replicates. The white dot shows median, the thick black line is the interquartile range (middle 50%), the thinner black line is the 95% confidence interval, and the colored area is the density plot. We note that the density plot is cut at the most extreme data points.

IsoCon’s precision increases with increased read depth, even when the isoform coverage is as high as 1,562x (Fig. S4). Such robustness is often hard to attain because increases in coverage beyond what is necessary for recall will only increase the number of errors in the data. The recall depends on the gene length and, hence, error rate. For *TSPY*, the recall becomes perfect at 17× coverage, while for *DAZ*, the recall reaches >90% at only 412x coverage (Fig S6). We expect accuracy to also be a function of gene copy similarity, *i.e*. a gene family that is generated using low mutation rate, thereby producing fewer variants between gene copies, has the potential to negatively affect IsoCon’s ability to separate transcripts. Somewhat surprisingly, accuracy decreases only slightly in these cases, and read depth has a much more substantial effect on accuracy than mutation rate or gene length.

Our experiments clearly indicate that IsoCon’s recall is strongly dependent on read depth. We investigated this in more detail by taking every transcript simulated as part of the experiments in Figure 1. We showed whether or not IsoCon captured a transcript as a function of the sequencing depth *(i.e*. total number of reads) of its respective experiment and its own sequencing depth (*i.e*. the number of reads that were sequenced from the isoform; Fig. 2). As expected, we see a strong correlation between IsoCon’s ability to capture an isoform and the isoform’s sequencing depth. For *TSPY*, IsoCon captures most transcripts with depth >3, while this number is approximately 10 for *HSFY*. Interestingly, the mutation rate plays only a minor role compared to the transcript depth. For example, for a fixed mutation rate of *DAZ*, there is a large range of transcript depths where some transcripts are captured and others are not. This indicates that other factors likely play an important role. We also observe that for *DAZ*, the minimum transcript depth required to capture an isoform increases as the total sequencing depth increases. This is likely due to the fact that the more reads cover an abundant transcript, the more reads are needed to recover a less abundant similar isoform. Another factor might be that the multi-alignment matrix becomes increasingly noisy as the number of sequences grows, negatively impacting both the error correction and the support calculation in the statistical test.

**Figure 2.**
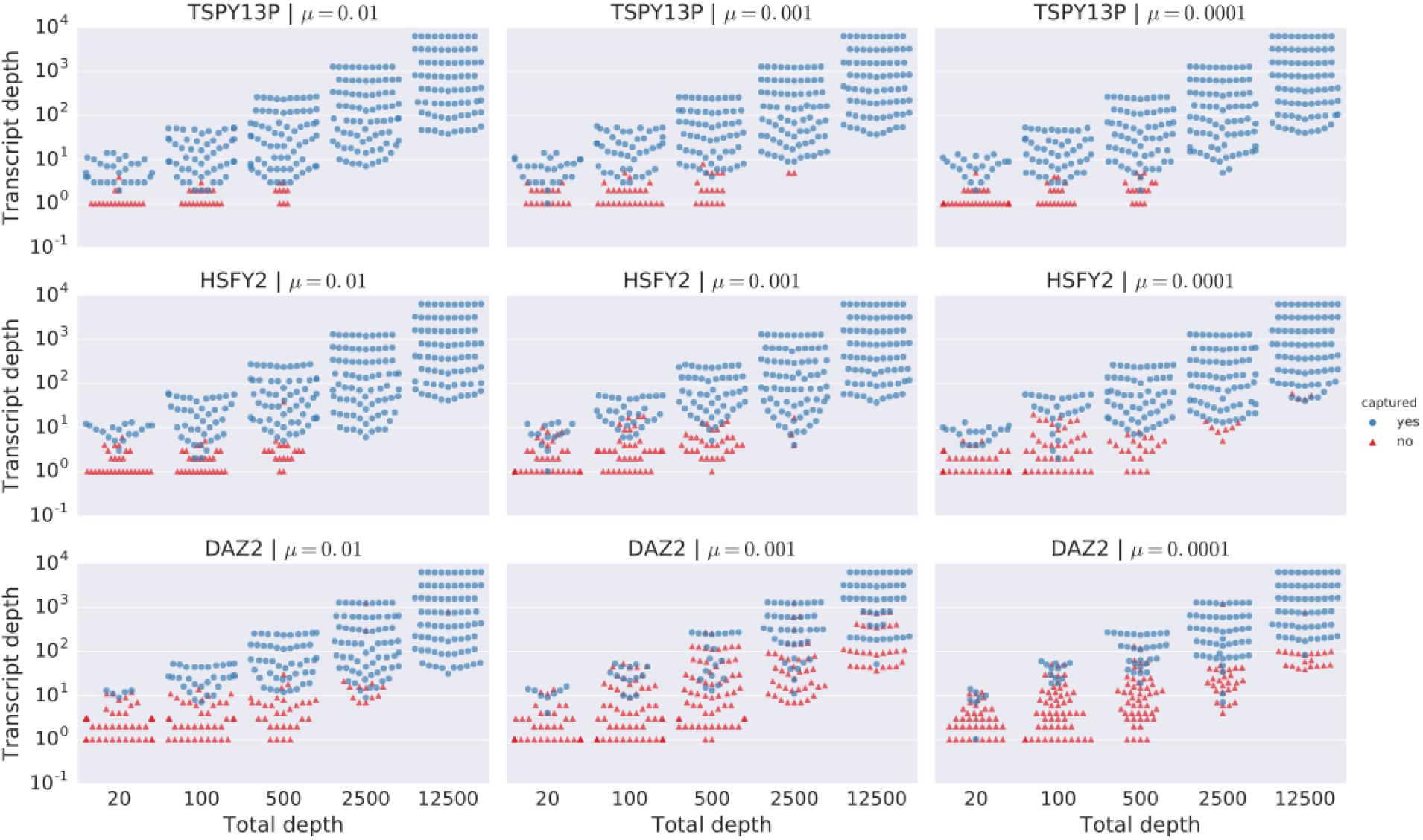
Power of IsoCon to separate transcripts with similar sequences as a function of coverage. Each isoform from the experiments in Figure 1 (including the ten simulated replicates) corresponds to a marker in this plot which is marked according to whether it is captured by IsoCon. This is a swarmplot generated with the seaborn package ^62^, which is a special kind of dotplot where the x-axis is categorical (total number of reads of the corresponding experiment) and points are spread out horizontally so as to not overlap each other. The y-axis shows the number of reads that were sequenced from the isoform, on a log scale. Isoforms that have no reads are not shown.

We also observe that IsoCon outperforms ICE in both precision and recall. ICE has poor precision which decreases with increased read depth. For example, at a read depth of 500 or higher, ICE’s precision is close to 0 in all our experiments. On the other hand, IsoCon’s precision is always >80% at read depths of 500 or higher. IsoCon also has higher recall in almost all cases for *HSFY* and *DAZ*. As for *TSPY*, the recall advantage fluctuates between the two algorithms but is fairly similar overall. Further investigation of IsoCon’s performance is detailed in Sup. Note C.

### Data from two human testes samples

As described in the experimental methods, we generated transcript sequencing data for nine ampliconic gene families for two human male testes samples using the Iso-Seq protocol from PacBio (Fig. S12 shows the number of passes per read) and, separately, using Illumina sequencing technology. We then used the ToFU pipeline ^17^ to filter out any PacBio CCS reads that either were chimeric or did not span a transcript from end to end. We refer to the resulting set as *the original* (CSS) reads. For the purposes of comparison, we ran IsoCon and ICE^17^ on the CCS reads. We also evaluated the proovread ^46^ tool, which uses Illumina reads to correct CCS reads (referred to as Illumina-corrected CCS reads). Supplementary Note D provides details on how these tools were run. We compared the results of the three approaches, as well as of the approach of just using the original CCS reads. Table 2 shows the number of reads generated and the number of transcripts called by each of these four different approaches.

**Table 2.**
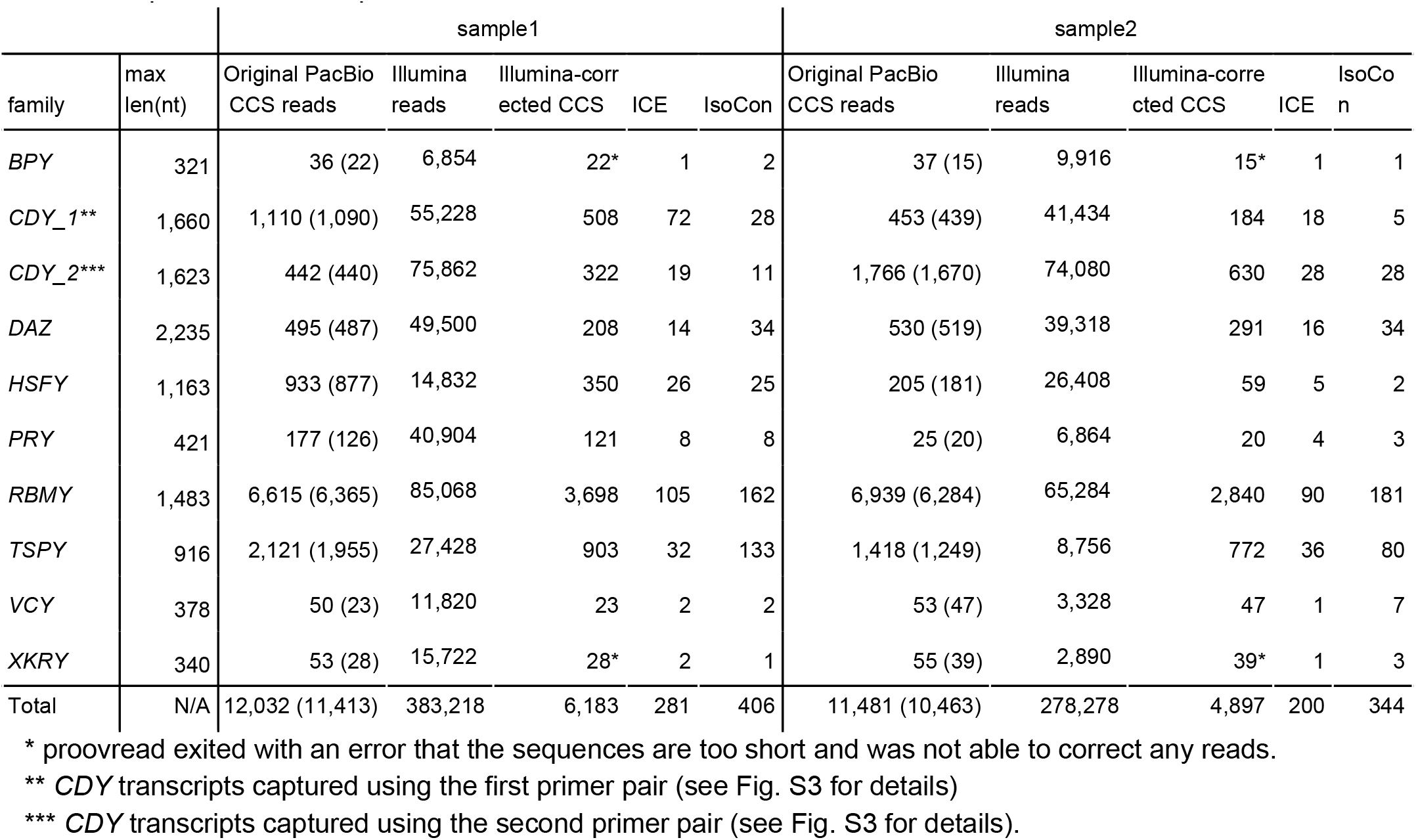
The number of reads and of predicted transcripts. We show: (1) maximum RT-PCR product length per family (including primers), (2) the number of original PacBio CCS reads, with the number of distinct read sequences in parenthesis, (3) the number of Illumina reads, (4) the number of distinct proovread Illumina-corrected CCS reads, (5) the number of ICE predicted transcripts, and 6) the number of IsoCon predicted transcripts. sample1 sample2 family max len(nt)

#### Validation

To validate IsoCon and compare its accuracy against other methods we used (1) Illumina reads, (2) internal consistency between samples, and (3) agreement with a transcript reference database.

We validated the nucleotide-level precision of IsoCon, ICE, Illumina-corrected CCS reads, and original reads with Illumina data generated for the same two individuals. Throughout all positions in the predicted transcripts, we classified a position as *supported* if it had at least two Illumina reads aligning to it with the same nucleotide as the transcript. Since Illumina sequencing depth was orders of magnitude higher than that of PacBio (Table 2), we expect most correct positions to be supported. Note that the lack of Illumina support does not always indicate an error, since Illumina’s GC-bias will result in some regions being unsequenced. However, we expect that the number of transcript errors is correlated with the number of unsupported positions. Figure 3 shows the percentages of supported nucleotides for each approach. On average, 99% of IsoCon transcript positions are supported, but only 93% of ICE transcript positions are supported. Similarly,96% of Illumina-corrected read positions and 79% of original read positions are supported. IsoCon has 70% of its transcripts fully supported (*i.e*. at every single position) by Illumina, compared to 2% for ICE, 15% for Illumina-corrected reads, and 20% for uncorrected reads.

**Figure 3.**
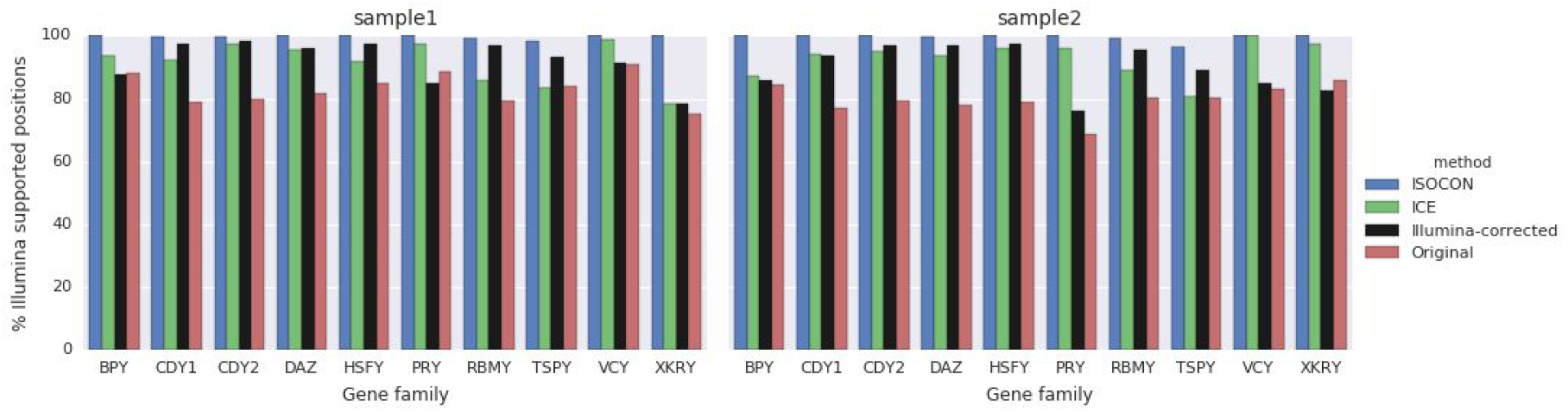
Percentages of positions in IsoCon/ICE/proovread, and original CCS flnc reads supported by at least two Illumina alignments with the same nucleotide.

While we expect some variability in the transcripts present in the two samples, we also expect a large fraction of them to be shared. IsoCon detected 121 transcripts that are present in both samples, corresponding to 32% of total transcripts being shared (averaged between two samples; Figure S9). The Illumina-corrected CCS reads shared 11%, while both ICE and the original reads shared less than 2%. This likely indicates the higher precision of IsoCon relative to other methods.

IsoCon also did a better job at recovering known Ensembl transcripts. We downloaded annotated coding sequences of the nine Y chromosome ampliconic gene families from the Ensembl database ^52^, containing 61 unique transcripts after removing redundancy (see Sup. Note E). We then identified database transcripts that were perfectly matched by the predicted transcripts. IsoCon had 21 matches to Ensembl, while ICE had only eight (included in those matched by IsoCon; Figure 4). IsoCon also had more matches to the database than the original reads, despite reducing the number of sequences by a factor of >29 (Table 2). Illumina-corrected CCS reads in total had one more exact match than IsoCon, but had >14 times more predicted transcripts to IsoCon, suggesting low precision.

**Figure 4.**
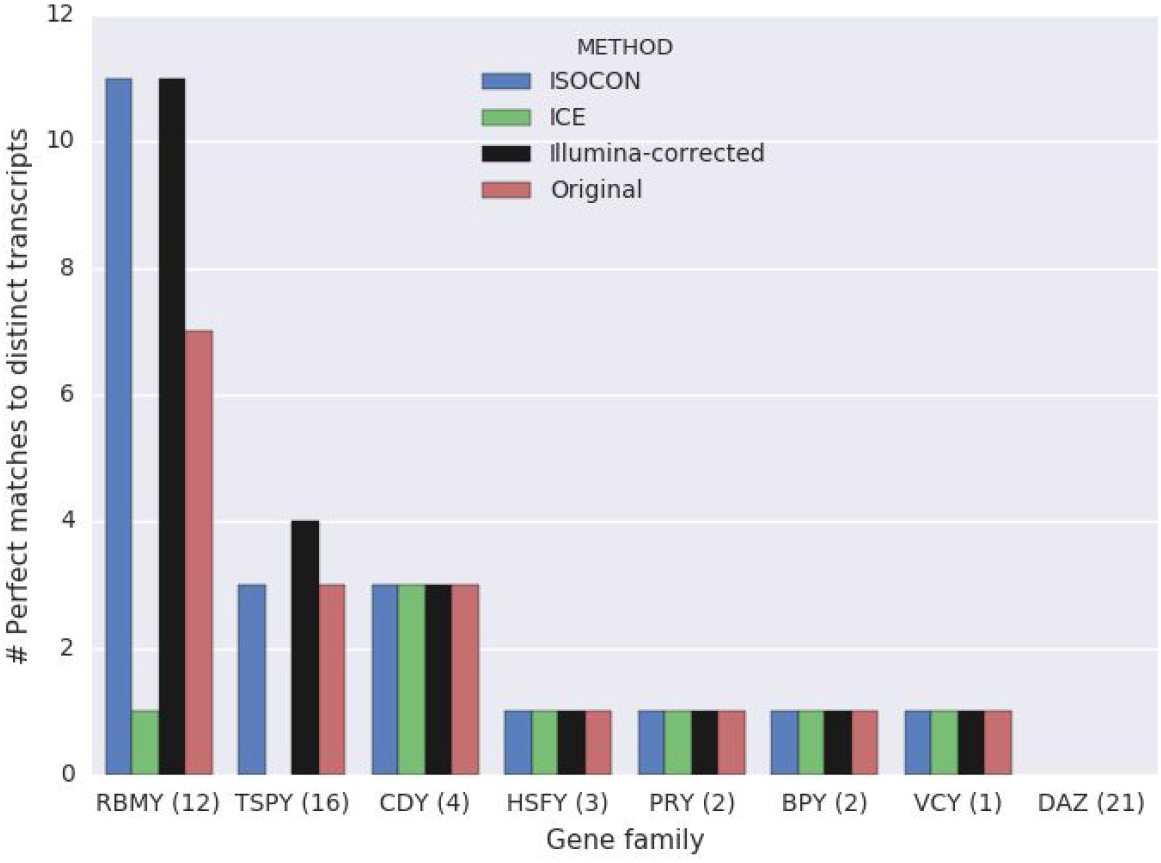
Ability of the methods to capture reference protein-coding transcripts in ENSEMBL database (no coding sequences are present for *XKRY*). The numbers in parentheses next to the gene family name in the x-axis indicate the number of unique transcripts in the database.

#### Isoform diversity

To study the transcripts IsoCon found, we first filtered out transcripts that were sample-specific. While we expect variability between the two individuals, such transcripts could also be false positives arising due to reverse transcriptase PCR (RT-PCR) errors. These errors, if present, were introduced prior to library construction and would be located in both Iso-Seq and Illumina reads. They would lead to unique sequences that would mimic true transcripts in both Iso-Seq and Illumina data. Since it is unlikely that identical RT-PCR errors would occur in two different samples, our downstream analysis only uses the 121 transcripts that were identically predicted by IsoCon in both samples. This also reduces any false positive IsoCon transcripts that might arise due to uncorrected sequencing errors. Table 3 shows the number of shared transcripts separated into gene families. We note, however, that the true number of transcripts in a sample might be higher due to sample-specific variants that we discarded. We further classified each transcript as protein-coding or non-coding depending on whether it is in-frame or out-of-frame with the human reference transcripts (see Sup. Note E for details). We found that 72 out of the 121 transcripts are coding, and five of the nine families harbor a total of 49 non-coding transcripts (Table 3), the other four families have only coding transcripts. We also found that 93 out of the 121 transcripts were not previously known, *i.e*. did not have a 100% match spanning the whole transcript when aligned to NCBI’s non-redundant nucleotide database (*nrint*). The multi-alignment for IsoCon’s *RBMY* transcripts -- the family with the most predicted transcripts -- is shown inFigure 5.

**Figure 5.**
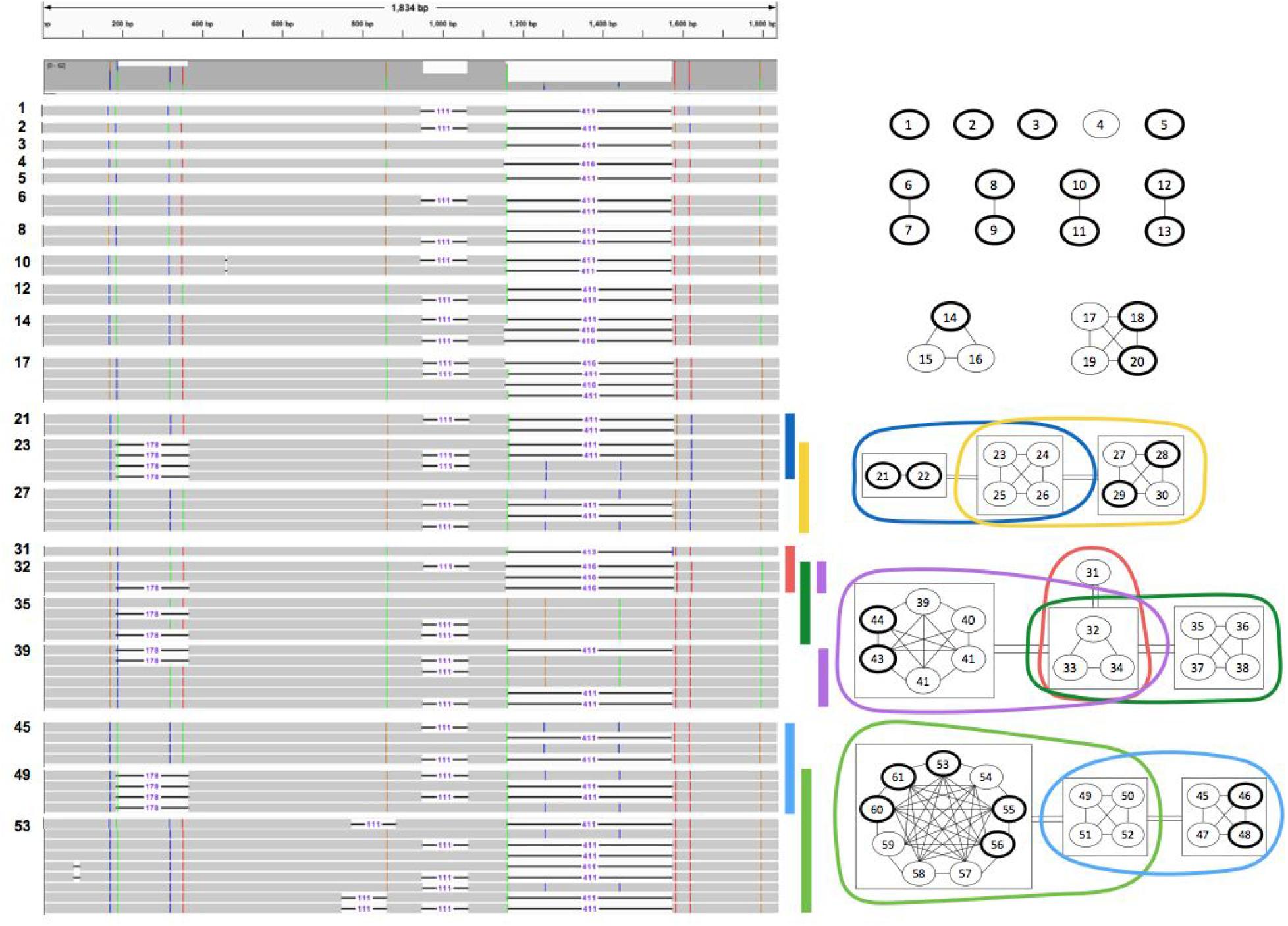
Illustration of the relationship between the 61 RBMY transcripts predicted by IsoCon and shared by both samples. Transcripts are indexed from 1 to 61. The left part of the figure uses IGV ^63,64^ to visualize a multiple-alignment of the transcripts. Colored positions are positions with variability in the transcripts while grey regions depict conserved positions. Deletions are shown with a horizontal line, with a number indicating their length. The right part of the figure illustrates the relationship between the 61 transcripts as a graph. Vertices are transcripts (labelled with their indices). A vertex is boldfaced if it is predicted to be protein-coding. An edge between two transcripts means that they are potential isoforms from the same gene copy, *i.e*., they have only exon presence/absence differences. To simplify the visualization, some of the vertices are surrounded by boxes, and a double-edge between two boxes indicates that all pairs of transcripts, between the two boxes, are potential isoforms from the same gene copy. Each maximal clique (*i.e*. *group* of vertices) greater than four vertices is shown as a colored circle. The colors of the circles correspond to the rows in the multiple-alignment that are marked with a similarly-colored vertical bar. A maximal clique should be interpreted as all transcripts that potentially originate from the same gene copy.

**Table 3.** IsoCon predicted transcripts shared between samples, separated by gene family and categorized as coding or non-coding. The calculated number of groups is shown, in comparison to known copy numbers from the reference genome and observed in human populations. Novel transcripts are those that do not have a perfect alignment to the NCBI transcript reference database, the numbers in parentheses indicate additional transcripts that have a perfect alignment only to ESTs, synthetic constructs, or *in silico* predicted transcripts.

| Gene family | Number of coding members annotated in the reference<sup>59</sup> | Range of copy numbers observed in human populations<sup>60,61</sup> | IsoCon transcripts shared between samples |  | Inferred group sizes |  | Novel transcripts |
| --- | --- | --- | --- | --- | --- | --- | --- |
|  |  |  | Num coding | Num non-coding | Number of groups (all) | Number of coding groups |  |
| *BPY* | 3 | 2-3 | 1 | 0 | 1 | 1 | 0 (0) |
| *CDY* | 4 | 2-4 | 3 | 2 | 2 | 2 | 1(1) |
| *DAZ* | 4 | 2-5 | 5 | 1 | 3 | **3** | 5(0) |
| *HSFY* | 2 | 2 | 0 | 0 | - | - | - |
| *PRY* | 2 | 2-3 | 1 | 2 | 1 | 1 | 1(0) |
| *RBMY* | 6 | 6-18 | 26 | 35 | 18 | 14 | 49(3) |
| *TSPY* | 35 | 12-38 | 34 | 9 | 20 | 14 | 37(3) |
| *VCY* | 2 | 2-3 | 1 | 0 | 1 | 1 | 0(0) |
| *XKRY* | 2 | 2-3 | 1* | 0 | 1 | 1 | 0(0) |
| Total | 32 | 32-79 | 72 | 49 | 51 | 38 | 93 |

\*The *XKRY* transcript shared by two samples for *XKRY*-based primers had a better alignment to *XKR3* than to *XKRY* and thus might not be Y-specific, but we did find a Y-specific transcript unique to sample 2.

#### IsoCon is sensitive to small and low-abundance variation

IsoCon was able to detect several transcripts even in the presence of an isoform with a much higher abundance that differed by as little as 1–3 bp. For example, one recovered IsoCon *RMBY* transcript in sample 2 was supported by only five reads and differed by only one nucleotide from a transcript that was supported by 863 reads. A second example is another IsoCon *RBMY* transcript that was supported by only five reads in sample 2 and differed by only one nucleotide from a transcript that was supported by 306 reads. Both of these lower-abundance transcripts were derived in both samples (the support for these transcripts in sample 1 was 10 and 9 reads, respectively), had perfect Illumina support, and were protein-coding. Neither of these were detected by ICE or present without errors in the original reads; however, both of them were also derived in the Illumina-corrected CCS reads. Figure 5 shows these two lower-abundance transcripts, indexed 3 and 1, respectively.

#### Separating transcripts into gene copies and comparing against known family sizes

A gene family consists of gene copies, each of which can generate several transcripts because of alternative splicing. In such cases, the transcripts will align to each other with large insertions/deletions (due to missing exons) but without substitutions. We determine the minimal number of groups (*i.e*. clusters) that are required so that each transcript can be assigned to at least one group and every pair of transcripts in the same group differs only by large insertions/deletions (see Sup. Note E for details). We refer to this as the *number of groups*, which is our best estimate of the number of copies, *i.e*. the size of each gene family. Note that allele-specific transcripts are non-existant for Y ampliconic gene families because of the haploid nature of the Y chromosome. We also determine the *number of coding groups*, which is the number of groups computed from the coding transcripts only. A group corresponds to the notion of a maximal clique from graph theory ^53^, and, since the number of transcripts is relatively small, the number of groups can be computed using a brute-force algorithm (Sup. Note E).

The number of groups for the nine different families are shown in Table 3, and Figure 5 illustrates the idea of groups using the *RBMY* family as an example. An important distinction between a group and a gene copy is that a transcript can belong to multiple groups. This happens if a transcript skips an exon that contains a variant separating two gene copies. In such a case, we cannot determine which copy it originates from, and our approach places it in both groups. The size of each group is therefore an upper bound estimate of the number of isoforms originating from each gene copy.

We note that the true number of copies in a gene family might be higher or lower than the number of groups determined by IsoCon, for several reasons. First, it is impossible to separate transcripts originating from copies with identical exonic sequences. As a result, we might underestimate the true number of copies. Second, copy number might differ between the two males analyzed ^54^. Because we are excluding transcripts unique to each male, we might underestimate the true copy number. Also, the copy number for a gene family might be the same between the two men, but some of the copies might have different sequences. Third, there may be copies that biologically differ only by the presence/absence of exons or by other large indels -- if there are no substitutions, the resulting transcripts would be grouped together by our approach. Since human Y chromosome ampliconic genes were formed by whole-region duplications, as opposed to retrotransposition ^55^, this situation should not be common. Nevertheless, if present, it would underestimate the number of copies. Fourth, RNA editing may generate transcripts that have substitutions but originate from the same copy, leading to overestimating the true copy number. Fifth, our approach to group transcripts relies on accuracy of transcript-to-transcript alignments, which can sometimes be inaccurate in the presence of repeats. With these caveats in mind, we nevertheless expect the number of groups to be a useful proxy for the size of the gene family.

We compared our number of coding groups against the number of copies annotated on the Y chromosome in the human reference genome (GRCh38/ hg38) and observed in previous studies of DNA variation in human populations (Table 3). For one of the gene families (*HSFY*), we did not find any transcripts shared between individuals. For four gene families (*CDY*, *DAZ*, *RBMY*, and *TSPY*), the number of coding groups falls within the previously observed range based on DNA analysis in human populations (Table 3). For the remaining four gene families (*BPY*, *PRY*, *VCY*, and *XKRY*), the number of coding groups is less than the copy number reported by prior studies. Three of these families -- *BPY, VCY* and *XKRY* -- had only one coding transcript shared between the two samples. Thus, overall, the number of coding groups is a conservative estimate of the number of ampliconic gene copies per gene family.

#### Running time and memory analysis

We compared runtime and memory of IsoCon, ICE, and proovread on our human samples (Table S2). We used a machine with an x86_64 system running Linux v3.2.0-4-amd64 and equipped with 32 2-threaded cores and 512 GB RAM. IsoCon is roughly 2x faster than ICE when running on a single core, and ∼5x faster when running over 64 processes. A direct comparison to proovread is challenging because it requires a minimum read length criteria that makes it unable to process the *BPY* and *XKRY* families. With this caveat in mind, proovread run times were 12% slower than IsoCon on 1 core and >3x slower on 64 cores.

## Discussion

We have used both simulated and experimental data to evaluate the performance of IsoCon and its alternatives. First, all approaches exhibit higher recall and precision compared to using uncorrected CCS reads. Second, our simulations uncovered the challenges that ICE has in deriving transcripts from high-identity gene families, as its precision deteriorates with increased read depth. On the other hand, IsoCon behaves as expected, with precision remaining fairly stable but recall reaching 100% with increasing read depth. On experimental data, our validations also confirm a low precision for ICE transcripts -- they are significantly less supported by Illumina data and are less consistent between samples, as compared with IsoCon. Similarly, the recall of ICE is lower than IsoCon’s as judged by matches to the Ensembl database. ICE transcripts only matched nine sequences from Ensembl, while IsoCon matched 22.

Third, for high-identity gene families, error correction using Illumina reads with proovread results in extremely low precision. This is possibly due to the challenge of multi-mapping reads, which arises when many similar long reads serve as alignment references. Nevertheless, Illumina-corrected CCS reads do recover an extra transcript from Ensemble, compared to IsoCon. In situations where low precision is tolerable to achieve a slightly improved recall, proovread may still be the preferred method.

IsoCon, similarly to ICE, uses an iterative cluster and consensus approach, but the two algorithms have fundamental differences. After clustering, IsoCon derives a weighted consensus based on the error profile within a partition, and uses it as information to error correct the reads; ICE, on the other hand, derives a cluster consensus using the stand alone consensus caller arrow, to be used in the next iteration without error correction of the reads. IsoCon and ICE also differ in the graphs they use to model the relationship between sequences and in the algorithm to partition the graph into clusters. IsoCon deterministically creates clusters modeled as a path-traversal problem, while ICE models a cluster as a maximal clique and uses a non-deterministic approximative maximal clique algorithm. Perhaps most importantly, IsoCon as opposed to ICE, includes a statistical framework that allows it to distinguish errors from true variants with high precision.

The experimental methods we use here have some potential limitations. First, very low-abundance transcripts might not be captured by Iso-Seq, since deep PacBio sequencing can be cost-prohibitive. Second, very short transcripts might not be captured during size selection. The size selection step of the PacBio library preparation protocol enriches for longer transcripts and thus some shorter Y ampliconic gene transcripts could not be captured. For example, we failed to retrieve some shorter *BPY* transcripts (Fig. S10). Both of these limitations can potentially be overcome via augmenting Iso-Seq data with Illumina RNA-seq data. Third, our approach of only sequencing the transcriptome does not provide a definitive answer to the size of each gene family and does not allow us to conclusively assign each transcript to a gene copy.

These limitations notwithstanding, IsoCon has allowed us to detect an unprecedented number of isoforms, as well as to derive better estimates on the number of gene copies in Y ampliconic gene families. Many of these isoforms are novel and need to be confirmed at the protein level. Nevertheless they are expected to open novel avenues of Y ampliconic genes research.

## Conclusions

We plan on improving IsoCon on several fronts as we continue to apply it to other gene families and species.

*Future Improvements to IsoCon: Computational*. IsoCon can be further improved by incorporating Illumina RNA-seq data (as opposed to just using it for validation, as we have done here). If IsoCon is adopted to cautiously make use of such data, it can further reduce false positives. Another avenue for improvement is to use data from the genome assembly or from known transcripts in the database; however, relying on such *a priori* information may be of limited use as we move towards deciphering transcripts from non-model organisms.

*Future Improvements to IsoCon: Experimental*. Several experimental improvements can be incorporated into IsoCon as well. First, to eliminate reverse transcriptase and sequencing errors, two replicate cDNA libraries can be prepared from the same sample ^56^. Any differences in transcript sequences between the two libraries will be consistent with such errors. Second, to identify sites with RNA-editing, one can amplify and sequence exons from the same individual, and compare RNA-and DNA-derived sequences. This will also assist in more conclusively determining the number and sequences of individual gene copies. Third, a PCR-based target enrichment approach, which we utilized here, might not have captured transcripts of gene copies with mutations at PCR primer sites. An alternative to this approach is to use a probe-based capture technique (e.g., ^57^), which does not depend on PCR primers.

*Future Improvements to IsoCon: Non-targeted approach*. Our ultimate goal is to develop a method that will allow one to decipher all the transcripts for a given sample and to apportion them to single-copy genes and individual copies of multigene families in the genome. A non-targeted Iso-Seq, *i.e*. sequencing of the whole transcriptome with PacBio technology, combined with IsoCon, can be employed to achieve this goal. A direct genome-wide application of Iso-Seq for this purpose is currently expensive and will result in predominant sequencing of highly abundant transcripts of housekeeping genes, leaving tissue-specific transcripts, frequently expressed at lower levels, undeciphered. Library normalization, applied to genome-wide, non-targeted transcripts, can overcome this limitation. While here we demonstrate an application of IsoCon to PacBio Iso-Seq data, in the future a similar approach can be applied to derive transcripts from direct sequencing of RNA with Oxford Nanopore technology ^58^.

## Acknowledgements

We would like to thank Qunhua Li for feedback on an earlier version of this algorithm. We would like to also thank Yale Center for Genome Analysis (YCGA) for PacBio sequencing service. This study was funded by National Science Foundation (NSF) awards DBI-ABI 0965596 (to K.D.M.), DBI-1356529 (to P.M.), IIS-1453527, IIS-1421908, and CCF-1439057 (to P.M.). This study was supported by the funds made available through the Penn State Clinical and Translational Sciences Institute, and through the Pennsylvania Department of Health (Tobacco Settlement Funds).

## SUPPLEMENTARY TABLES & FIGURES

**Table S1 (SupTableS1.csv).** RT-PCR primers designed in the first and last coding exons of the nine Y ampliconic gene families. Each primer starts with a 21 bp-long PacBio barcode that is unique for each sample.

**Table S2.**
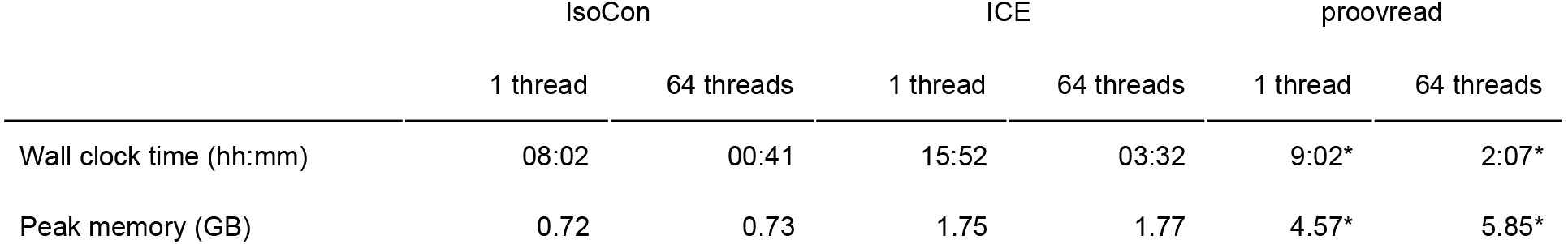
Runtime and peak memory usage of IsoCon, ICE, and proovread to process the whole biological dataset. Note that proovread did not perform any correction of reads from the *BPY* and *XKRY* families, due to their short size.

**Figure S1.**
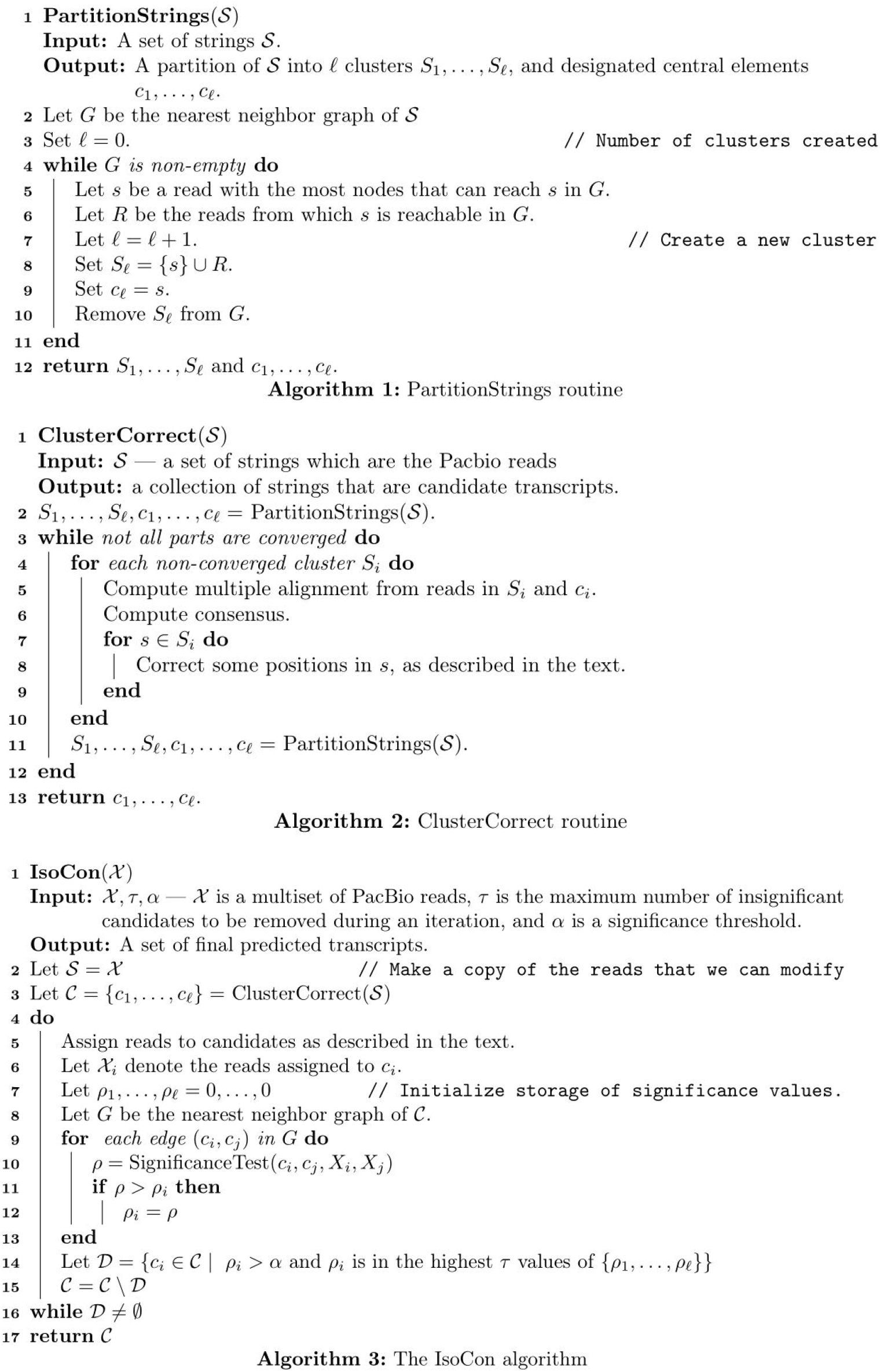
Pseudo-code for IsoCon algorithm.

**Figure S2.**
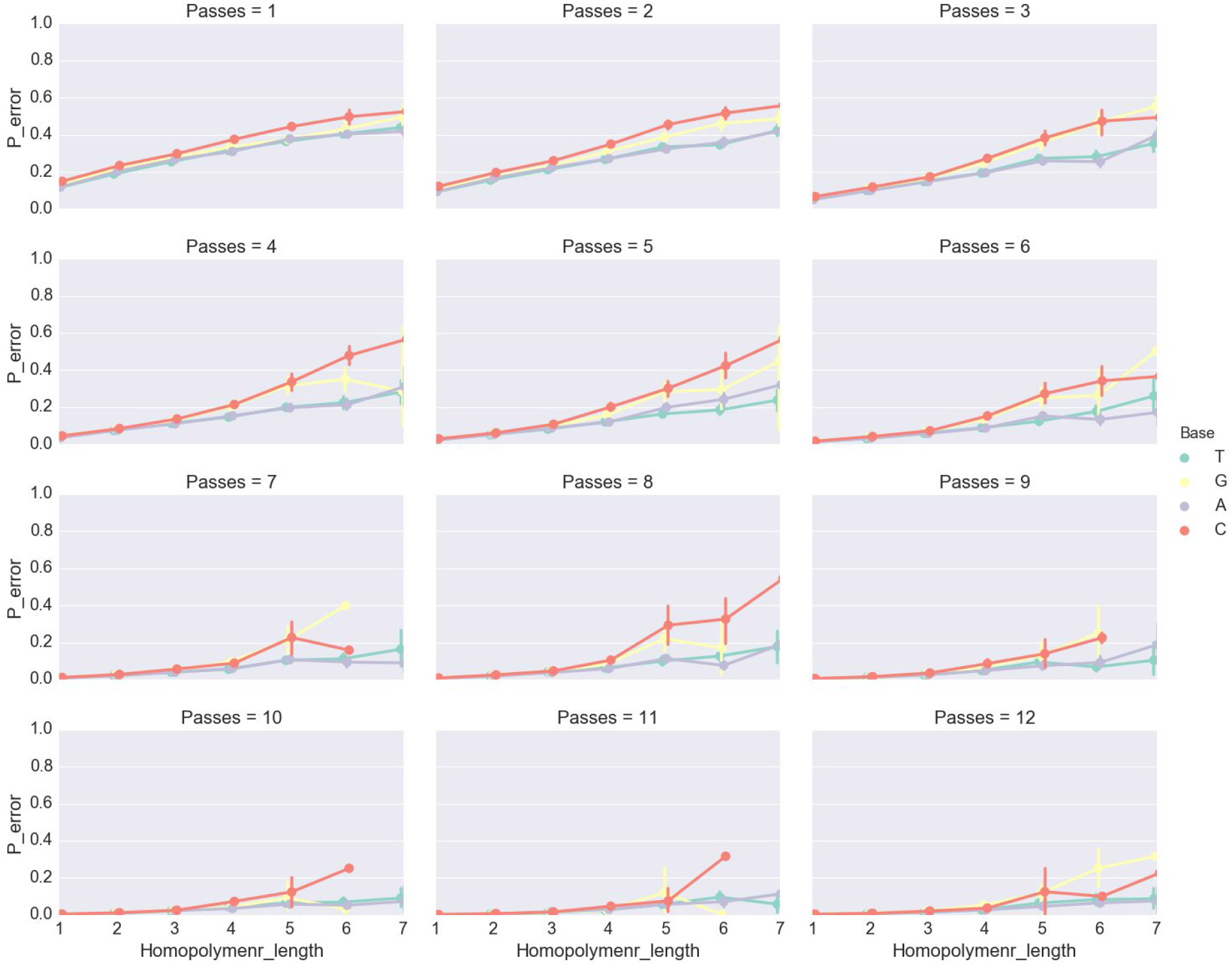
Quality predictions from PacBio’s base calling algorithm Arrow 65 for CCS reads based on a subsample of 5,000 CCS reads. Each panel shows Arrow’s prediction that the called base is wrong (y-axis) as a function of the homopolymer length in a CCS read (x-axis). The predictions are split by the four different bases. Each point within a line is the mean probability of error for a given nucleotide and the length of the homopolymer that contains it. Vertical lines for each point are the 95% confidence interval of the mean. Each of the panels correspond to the shown number of passes over the read.

**Figure S3.**
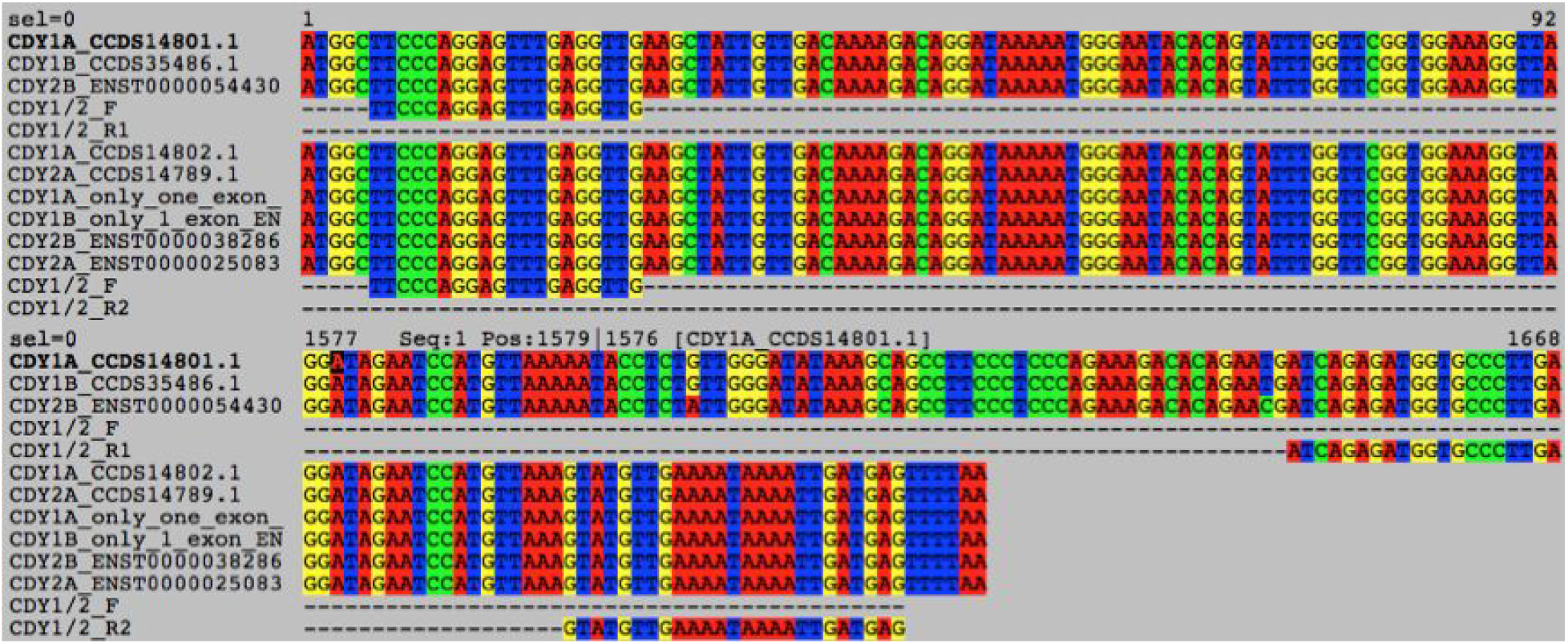
Alignment of two *CDY* primer pairs (CDY1/2_F and CDY1/2_R1;CDY1/2_F and CDY1/2_R2) to transcripts from Ensembl database showing the necessity of designing two alternative reverse primers to capture all protein-coding transcripts for this ampliconic gene family.

**Figure S4.**
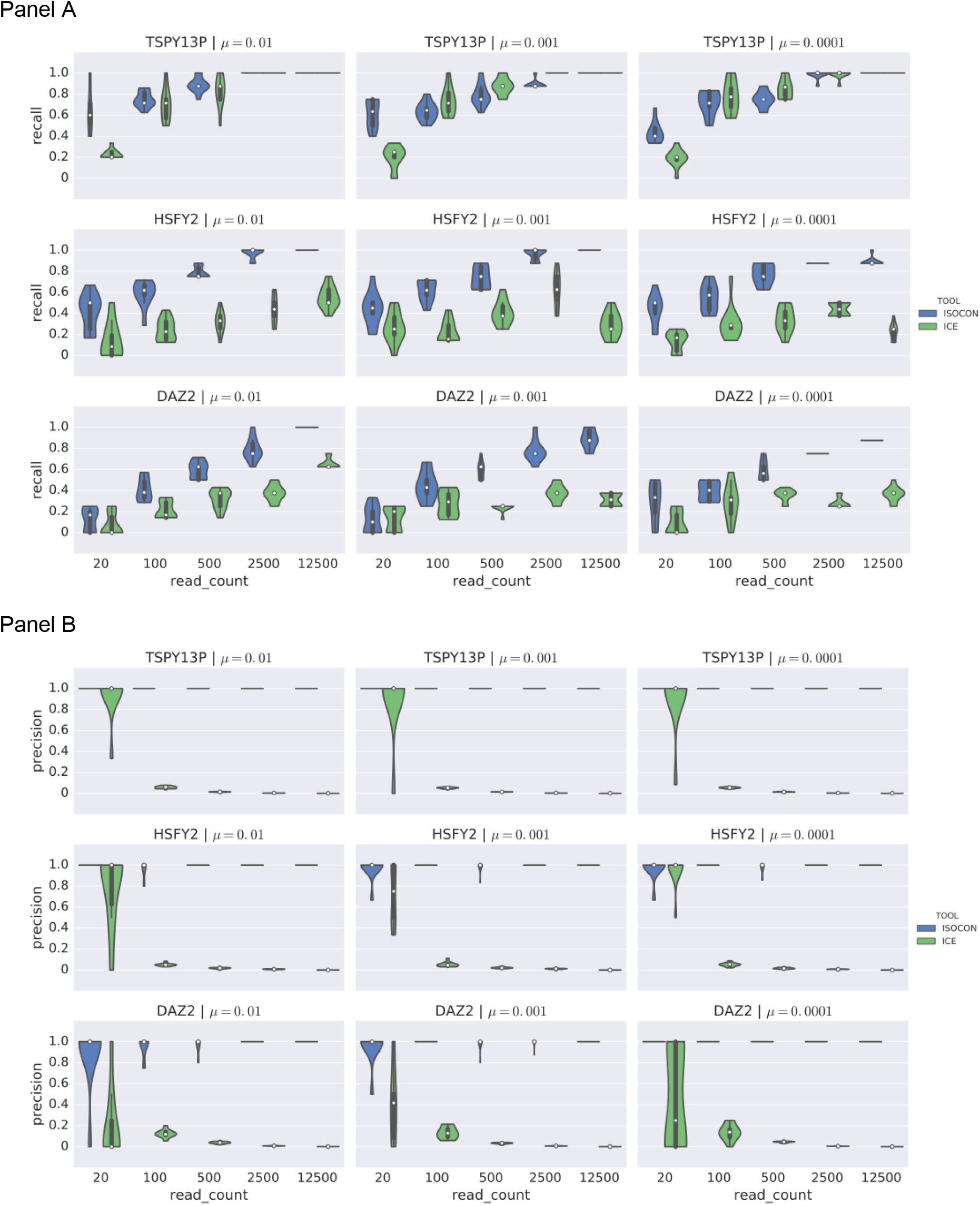
Violin plots showing the recall (panel A) and precision (panel B) of IsoCon and ICE on simulated families of transcripts with the same exon structure and unequal abundance rates. Each plot shows results for a total of 8 isoforms with abundances randomly assigned and ranging from 0.4% to 50%.

**Figure S5.**
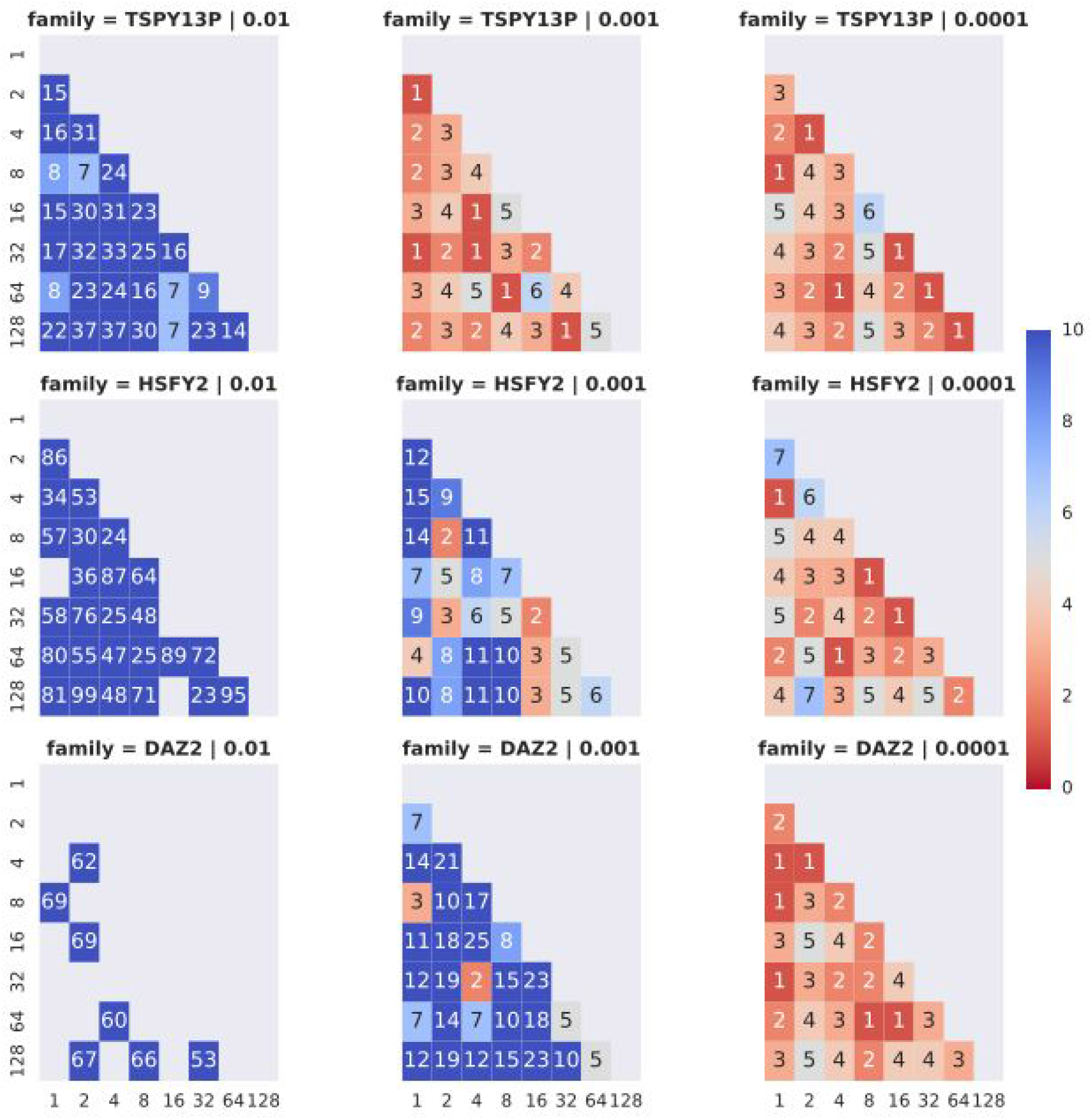
Edit distance between gene copies for simulated data sets of 8 gene copies. Each matrix shows edit distances between the 8 simulated copies for a specific gene family and mutation rate. The x and y axis show the abundance level of the copy in the unequal abundance experiment. The number in each block shows the edit distance between copies. Numbers on or above the diagonal are masked. Also masked are any blocks with edit distance >99.

**Figure S6.**
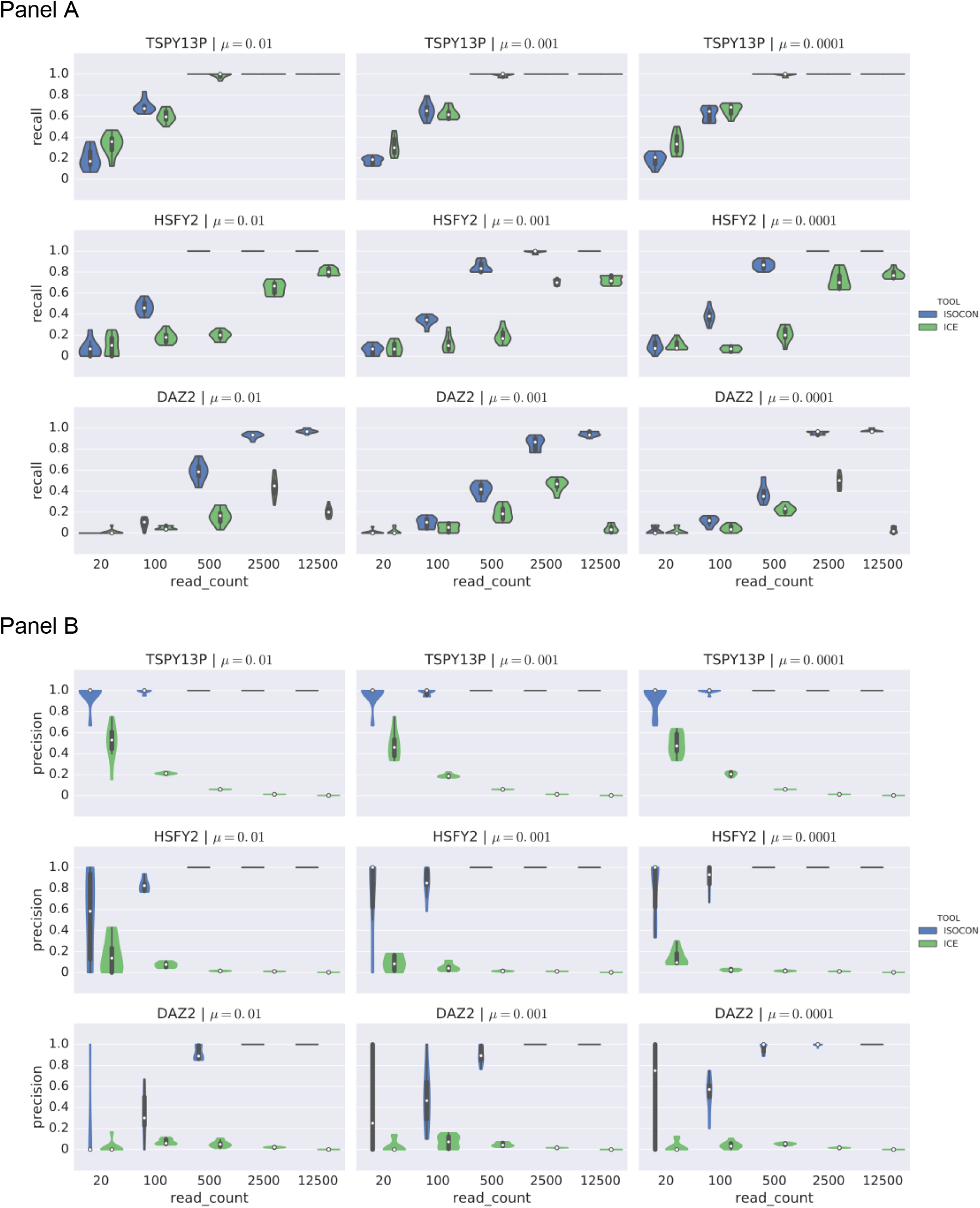
Violin plots showing the recall (panel A) and precision (panel B) of IsoCon and ICE on simulated families of transcripts with different exon structure and equal abundance rates. Each plot shows results for a total of 30 isoforms with equal abundances.

**Figure S7.**
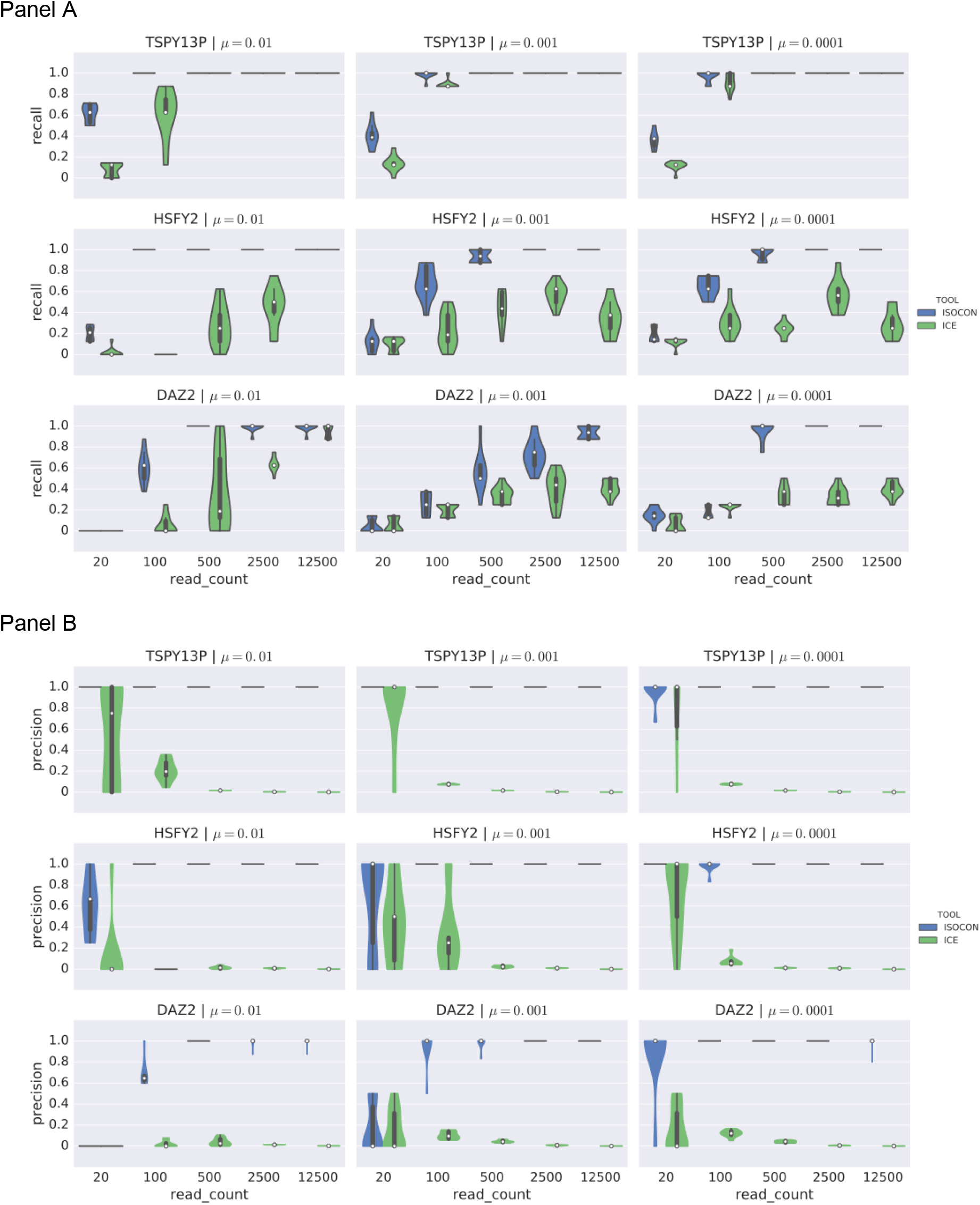
Violin plots showing the recall (panel A) and precision (panel B) of IsoCon and ICE on simulated families of transcripts with the same exon structure and equal abundance rates. Each plot shows results for a total of 8 isoforms.

**Figure S8.**
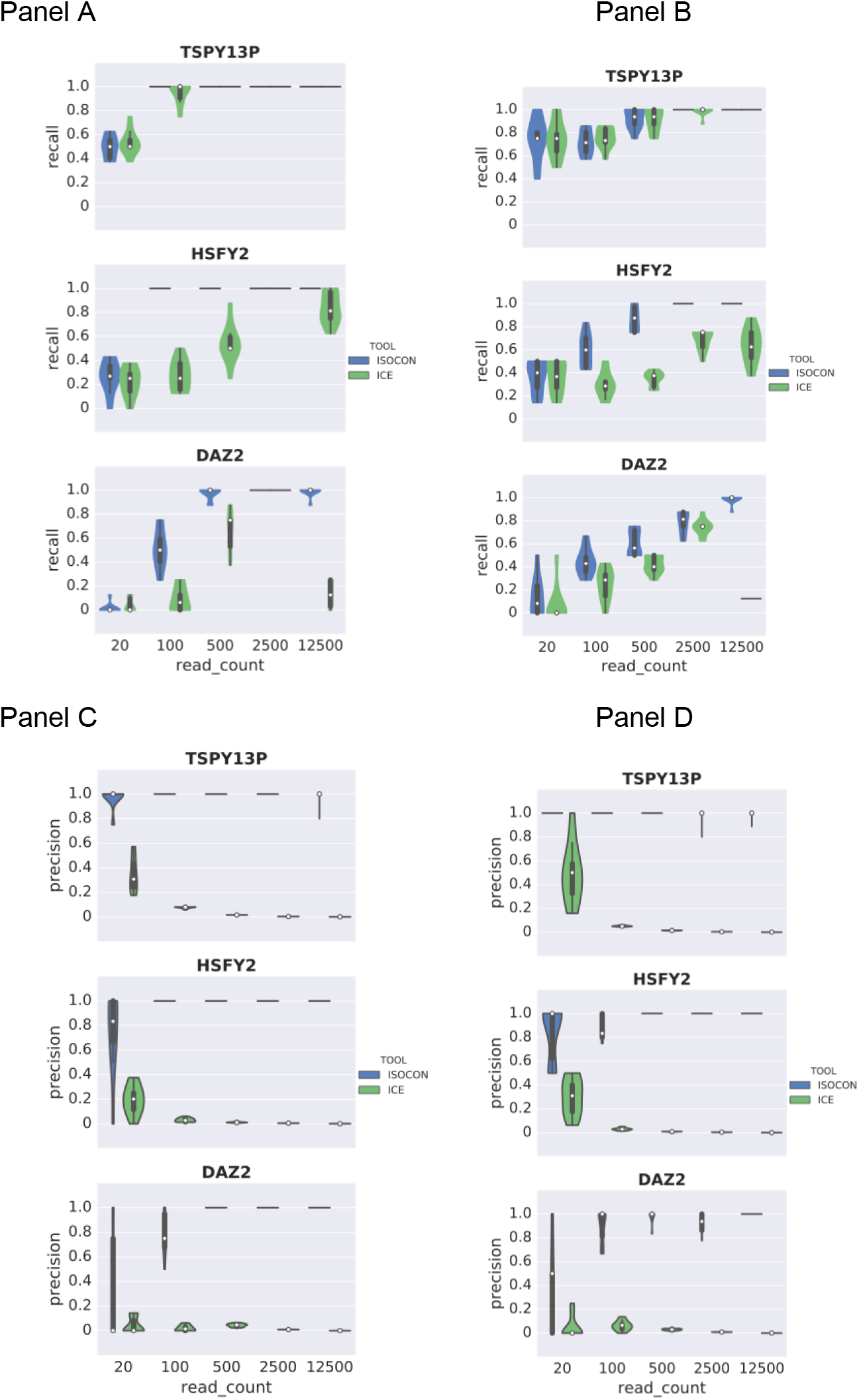
Violin plots showing the recall (panel A-B) and precision (panel C-D) of IsoCon and ICE on a single gene copy with eight different isoforms with equal abundance rates (left) and unequal abundance rates (right).

**Figure S9.**
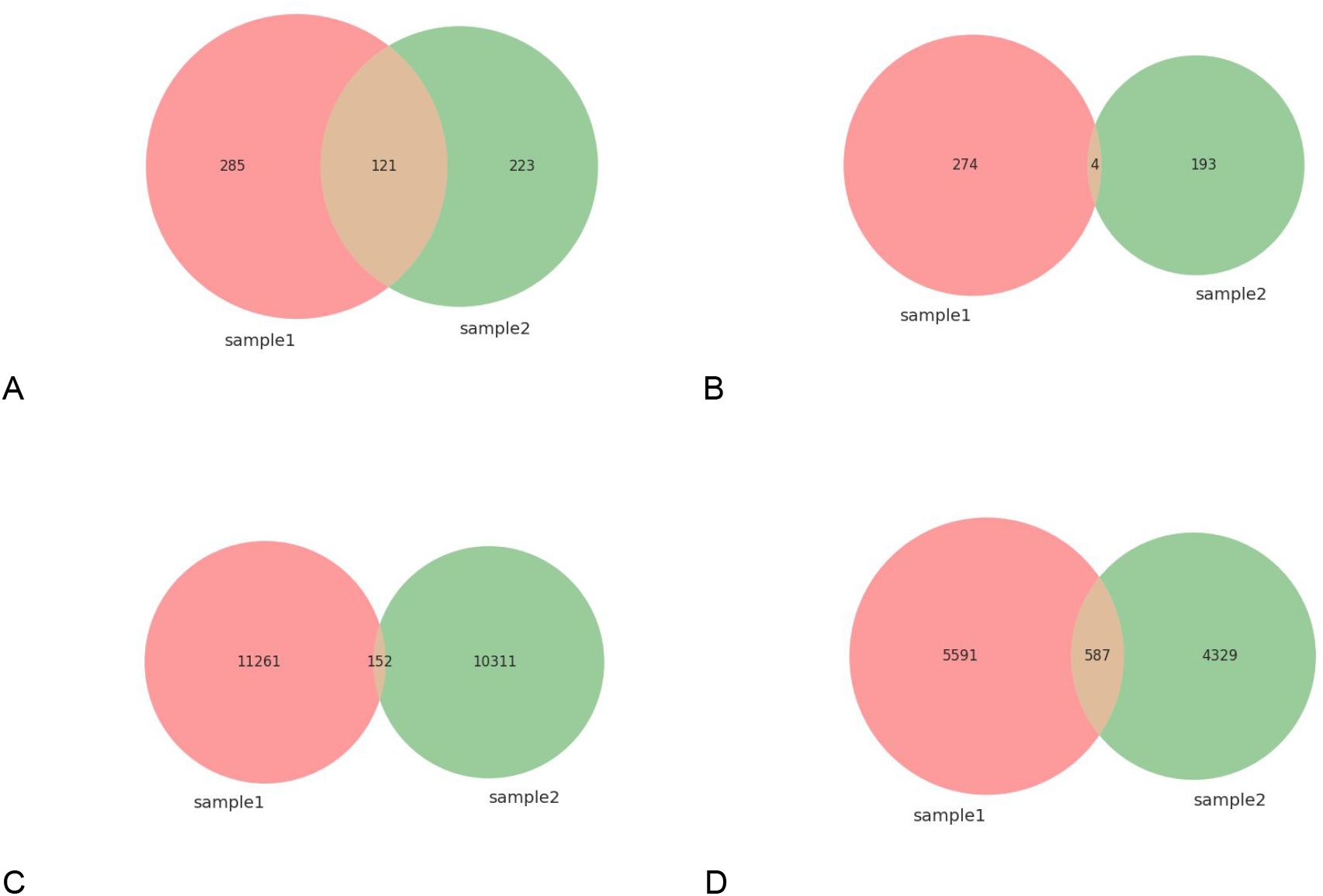
Venn diagram of the number of predicted transcripts shared between two samples for (A) IsoCon, (B) ICE, (C) original reads, and (D) Illumina corrected CCS reads. A transcript is shared if it has a perfect match between samples (edit distance of 0). Identical sequences within one sample are collapsed.

**Figure S10.**
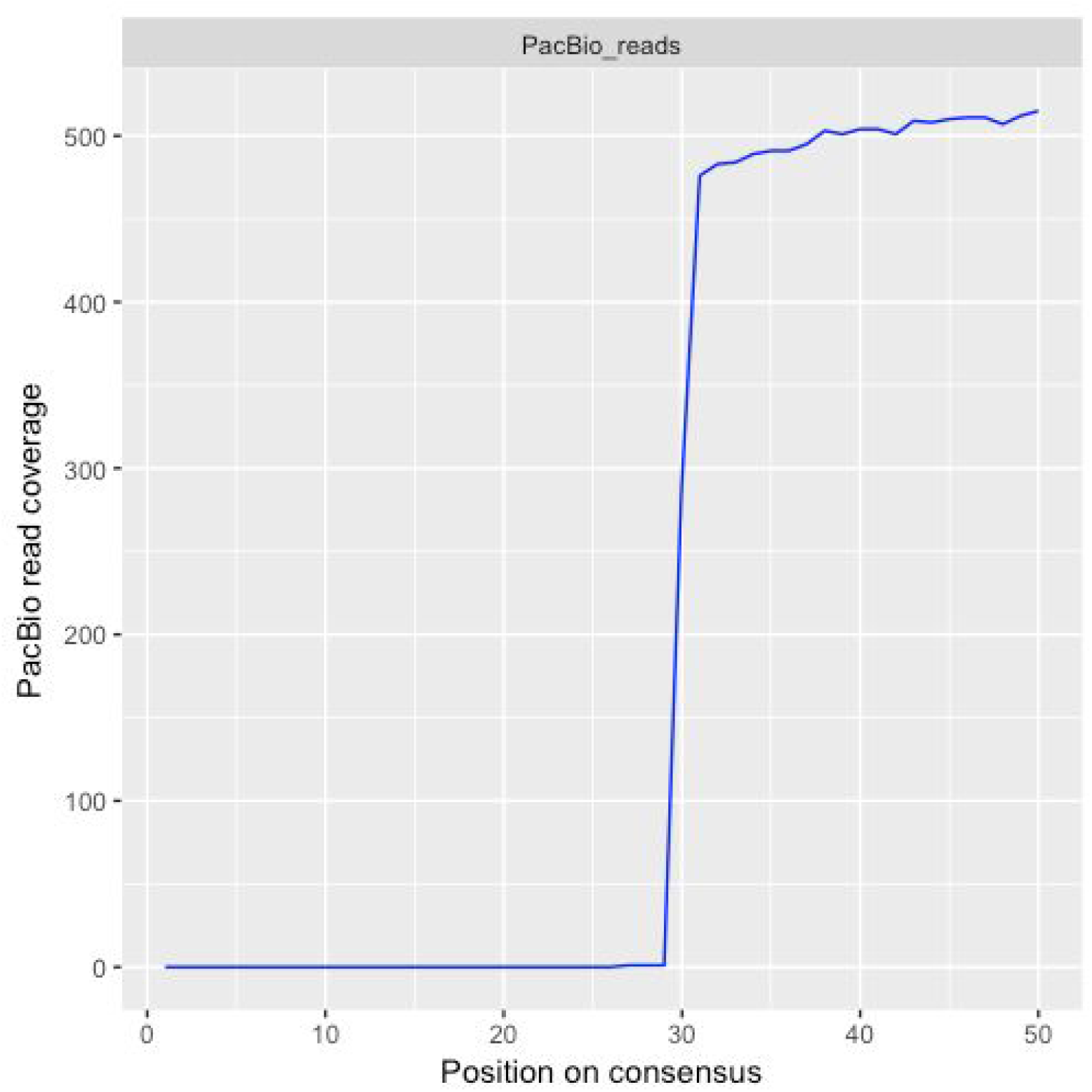
Coverage of CCS reads along the first 50 bp of the *BPY* consensus. To have the most complete set of reads, we combined the original CCS reads together with those that did not completely span a transcript from end to end. None of the reads includes the first 26 bp from the 5' end (including the 17 bp-long forward primer). This might be due to a limitation on the fragment size for library construction for PacBio sequencing.

**Figure S11.**
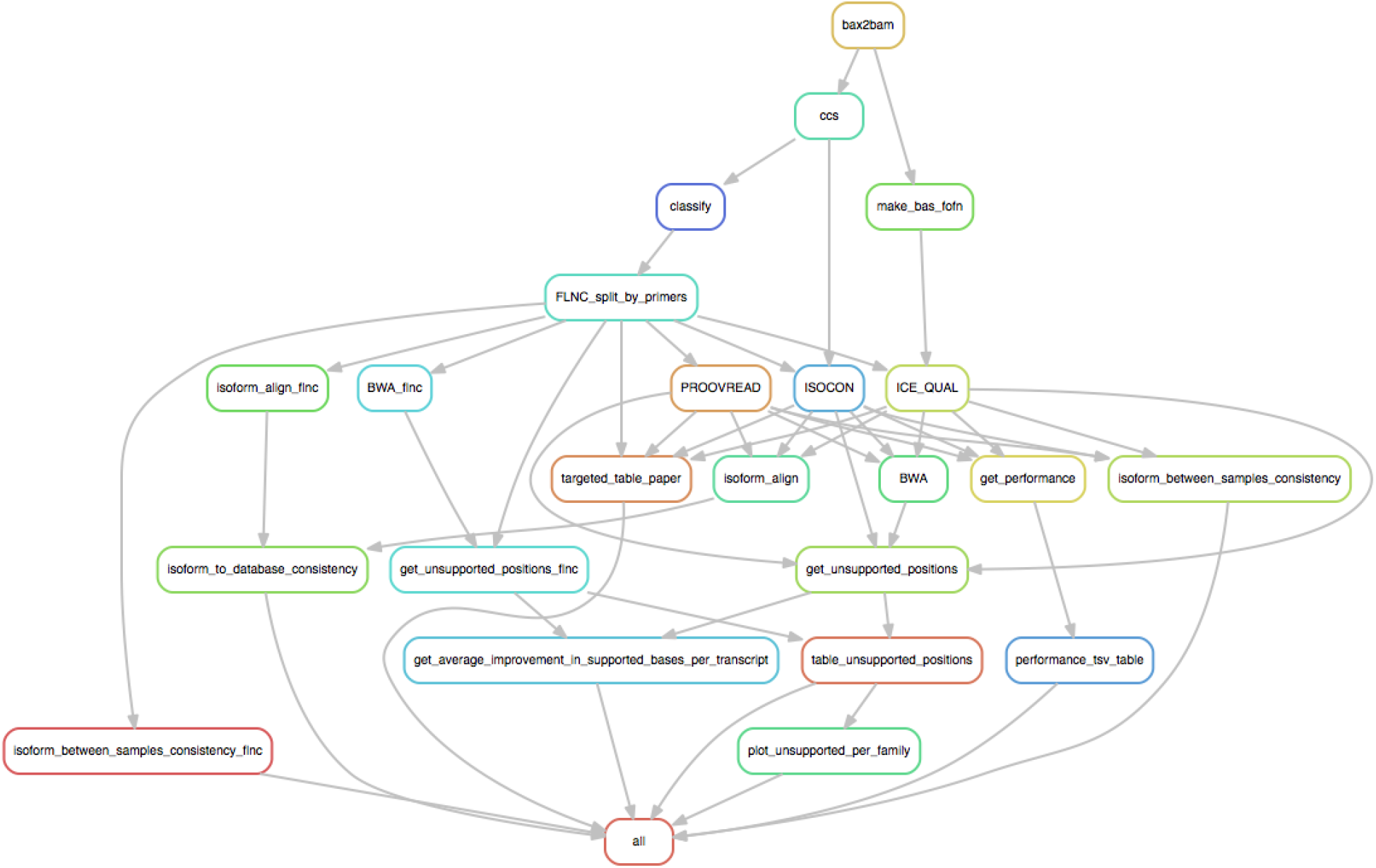
Biological data processing workflow with snakemake. bax2bam, ccs, and classify are tools included in the PacBio smrtlink v4.0 tool suite. The rule “split_by_primers” splits the reads into batches for each given primer.

**Figure S12.**
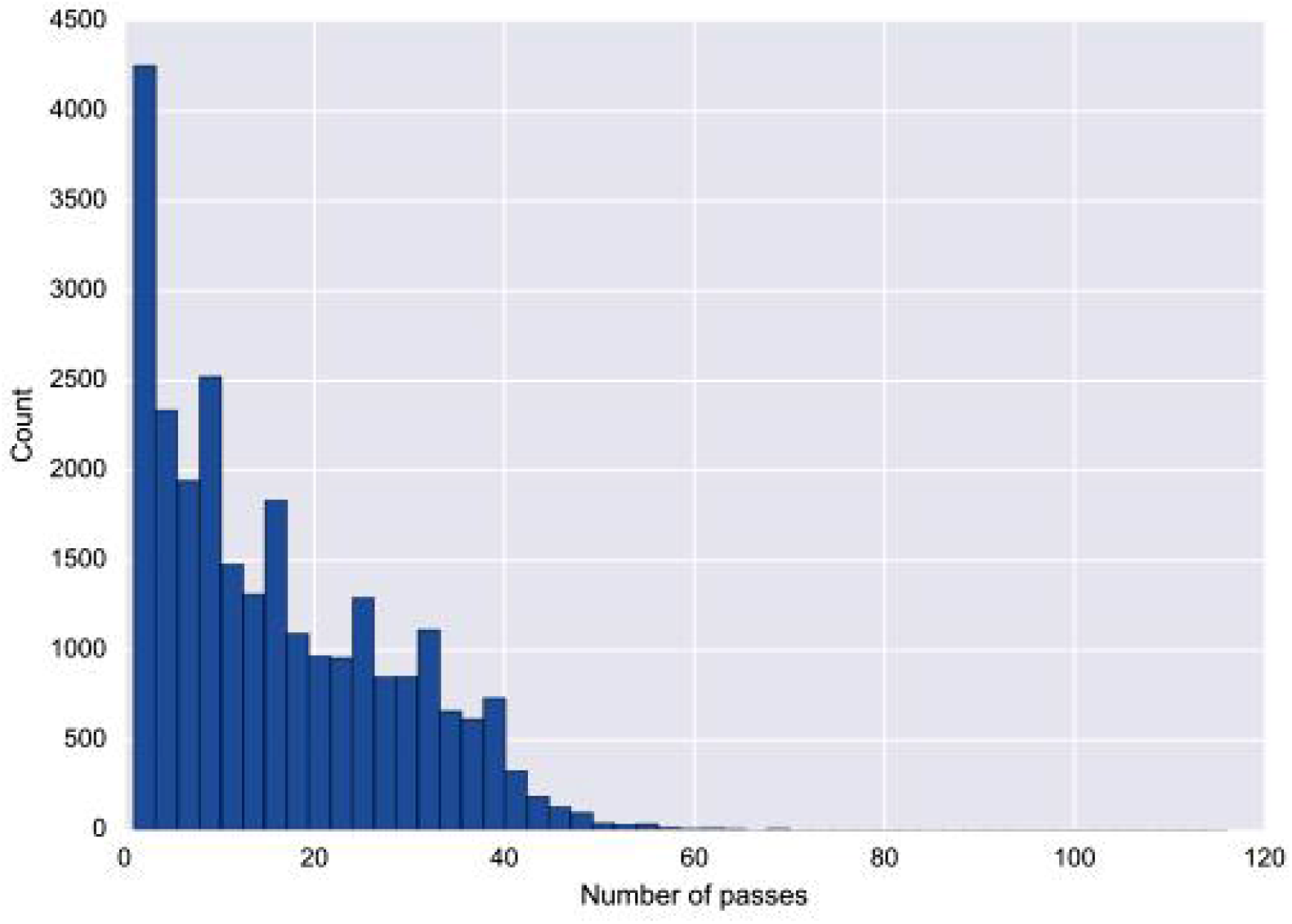
Histogram showing the number of polymerase passes in the CCS reads for both samples. The average number of passes is 16 and the median is 13.

